# Suboptimal human inference reflects an efficient and flexible information bottleneck

**DOI:** 10.64898/2026.06.10.731461

**Authors:** Jacob A. Parker, Alexandre L.S. Filipowicz, Kristen Li, Vijay Balasubramanian, Joseph W. Kable, Joshua I. Gold

## Abstract

Human inference is often suboptimal in ways that vary across individuals and tasks. We propose that at least some of this variability reflects information-processing limits. To test this idea, we developed a task-general application of the information-bottleneck framework to quantify how much information individuals use (information capacity) and how effectively they use it (information efficiency). We applied this framework to human choice behavior on two distinct inference tasks and found that individual differences in performance were driven largely by variation in information capacity, while information efficiency remained consistently high. This pattern of variable capacity and high efficiency also applied to a subset of participants who appeared inefficient overall, but were efficient when evaluated relative to a simplified (heuristic) encoding of the observations. Regardless of whether participants based their strategies on full or simplified encodings, they tended to adjust their performance to different task demands by changing their information capacity while maintaining efficiency. We interpret these findings in terms of new analytical results linking optimal inference under information-capacity constraints to evidence-based choice noise, suggesting that a key driver of human behavioral variability is rational adaptation to flexible information-processing limits.

## 1 Introduction

Our decisions often depend on latent features of the environment that cannot be observed directly but instead must be inferred from what we observe. For example, one might infer that it will rain soon based on observations about cloudiness, windiness, temperature, and humidity, then use that inference to decide to seek shelter. For tasks requiring such inferences, human decision-making behavior varies widely across individuals. This variability often manifests as a broad range of seemingly suboptimal inference strategies that differ in their degree of departure from optimal Bayesian inference [1–4]. This high degree of individual variability poses a substantial challenge to elucidating general principles that govern inference in the brain.

A possible explanation for this wide range of suboptimal behaviors is bounded (or resource) rationality [5–7]. That is, decision-makers strive to perform well, but to do so they must make use of limited and costly resources such as time, energy, and computational capacity. Balancing the competing needs of maximizing performance and minimizing costs leads to a range of inference strategies, from those that are maximally accurate but resource intensive, to those that are resource-frugal but potentially (but not always) less accurate. Bounded rationality redefines optimal choice behavior as achieving the highest possible performance given the resources used. From this perspective, a key to understanding human decision-making is to identify the specific resources whose limited use govern inference.

One source of limited resources identified in previous studies relates to the computational costs associated with algorithmic complexity. For example, formalisms of computer science and related fields have been used to quantify the different costs associated with a given algorithm in terms of which and how many mathematical and memory operations are used, how long it takes, and other factors [2, 8–12]. These results can then be used to provide insights into how humans balance computational complexity with performance [2]. However, one drawback to these approaches is that they require making strong assumptions about the exact algorithm(s) the brain is using to perform a particular task, which generally cannot be known with certainty.

Here we bypass such assumptions by proposing a form of bounded rationality that focuses on information use, agnostic to the algorithmic form of the strategy used to process that information. Specifically, under the premise that it is costly to encode and use information to form inferences, we propose that people flexibly adjust the information capacity of their inference processes. We quantify this capacity, and its impact on performance, via the information bottleneck (IB) [13]. This information-theoretic framework, which is related to rate-distortion theory [14], has been used to show how certain biological and artificial neural systems balance limits in information capacity with performance [15, 16]. In a novel application of this framework, we show that it can be applied directly to choice behavior of individual participants performing simple inference tasks. Our findings shed new light on the nature of bounded rationality that governs how we make inferences about the world.

## 2 Results

We applied the IB framework to human choice behavior on two classic inference tasks (Fig. 1), both conducted using the Prolific online platform. The “bead-prediction task” [17] (based partly on preregistration doi:10.17605/OSF.IO/KZXNQ; see Methods for details) required participants to predict the color of the next bead drawn from one of two hidden source jars that contained different ratios of black and white beads and could switch probabilistically between trials as the source of the draws (Fig. 1A). The “horse-prediction task” (a form of weather-prediction task [4, 18, 19]; preregistration: doi:10.17605/OSF.IO/HQWCY) required participants to predict which of two horses (latent states) would win a race based on a combination of shape cards (observations) shown on each trial (Fig. 1B). Thus, both tasks required inferences about which of two latent states generated a set of observations, but based on different kinds of observations with different probabilistic structures. Below we first derive properties of the IB as it relates to these kinds of tasks, then apply those findings to the analysis of behavioral variability across and within participants for the two tasks.

**Fig. 1:**
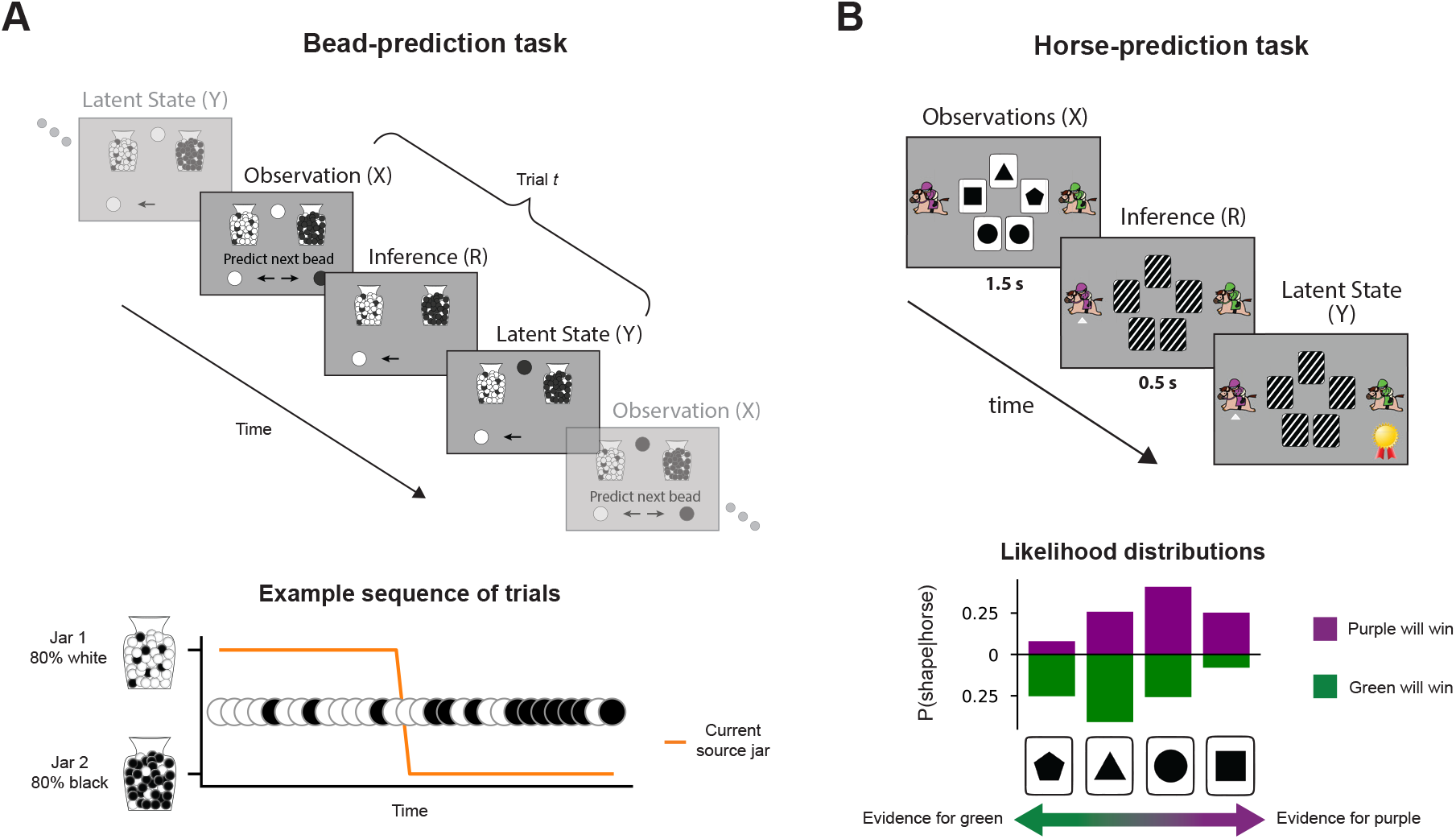
Inference tasks. **(A)** Bead-prediction task. On each trial, a bead was drawn probabilistically from one of two hidden source jars containing different mixtures of black and white beads. Participants predicted the color of the bead that would be drawn on the next trial, requiring an inference about the current jar. The source jar switched from one trial to the next with a low probability (hazard rate), implying that accurate inferences should depend on integrating sequences of bead draws over multiple trials. **(B)** Horse-prediction task. On each trial, five shape cards were sampled independently with replacement from one of two likelihood distributions, corresponding to the eventual winning horse. Participants inferred the winning horse based on these sampled cards, whose evidence strength was characterized by log-likelihood ratios that could be summed together to perform optimal inference.

### 2.1 IB framework

In general, the IB framework [13] can be used to assess the following question: if some signal *X*, which contains information about another signal *Y*, is compressed, how much information does the compressed signal retain about *Y* relative to the optimal compression? In our application to human inference (Fig. 2A), *x* ∈ *X* are the discrete observations (e.g., combinations of shape cards for the horse-prediction task) used to infer (or predict) *y* ∈ *Y*, which is an unobservable “latent state” (e.g., which horse will win). Now let *r* ∈ *R* be the potentially noisy inference report (e.g., prediction of which horse will win) made about *Y* after observing *x*. This *r* is effectively a compressed version of *x* that can, in principle, retain all the information *x* contains about *Y*. The inference strategy is characterized by *p*(*r*|*x*), the evidence-dependent choice probabilities.

**Fig. 2:**
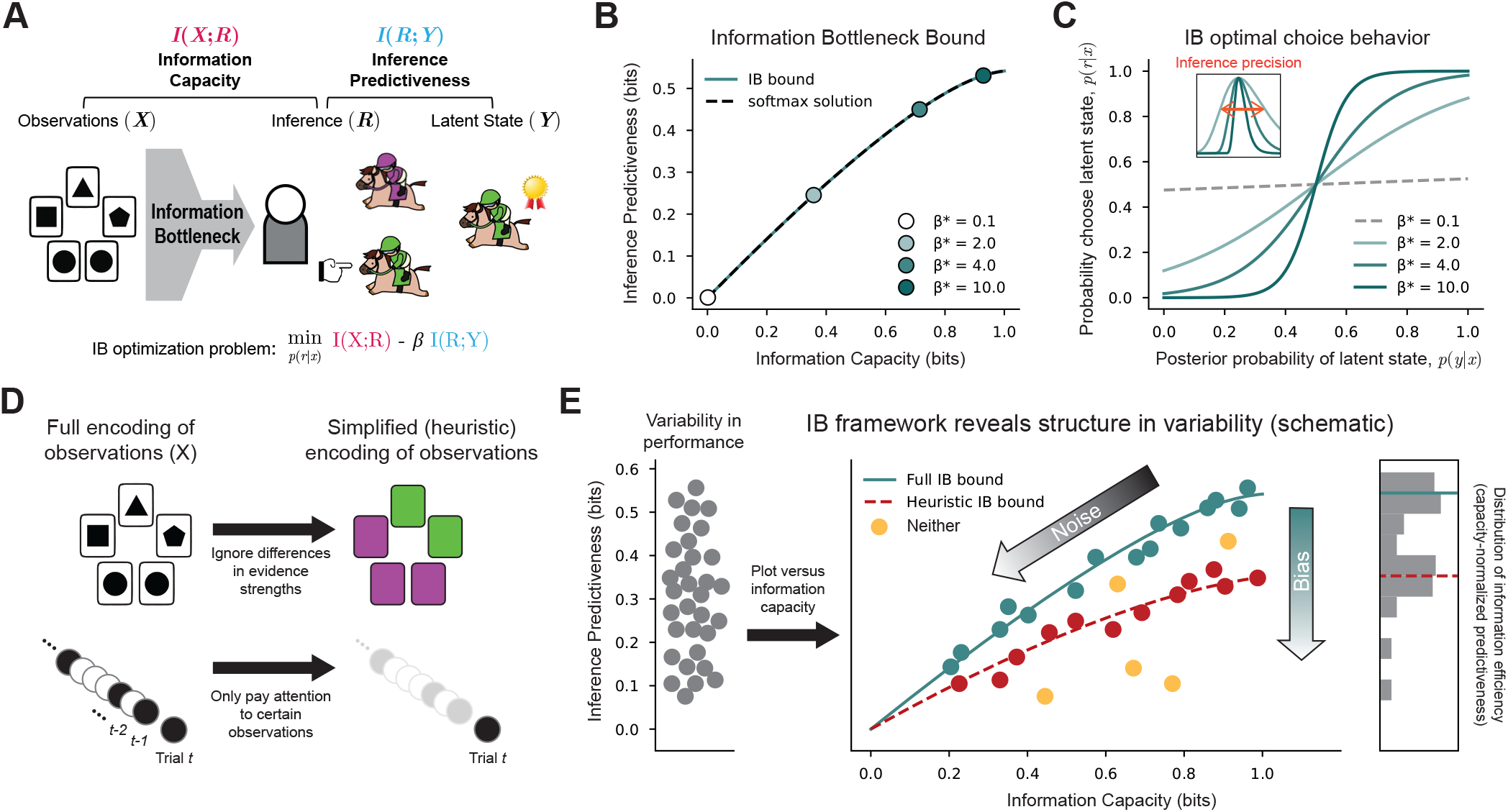
Information-bottleneck (IB) framework. **(A)** The IB framework formalizes the tradeoff between: 1) minimizing information-encoding and processing costs, which we formalize as the mutual information between the observations *X* and inferences *R*, *I*(*X*; *R*) (“information capacity”); and 2) maximizing accuracy, which we formalize as the mutual information between *R* and the latent states *Y* one is trying to infer, *I*(*R*; *Y*) (“inference predictiveness”). The relative strength of these objectives is controlled by *β*. **(B)** Solving the IB optimization problem (here for a variant of the horse-prediction task) over many values of *β* yields pairs of *I*(*X*; *R*) and *I*(*R*; *Y*) corresponding to the IB bound (solid line), which gives the maximum possible predictiveness (y-axis) as a function of information capacity (x-axis). Under certain conditions (see text and Math Appendix), the solution simplifies to a standard softmax form. Varying the inverse temperature parameter *β*^∗^ (colored points) corresponds to solutions along the IB bound (dashed line). **(C)** Psychometric functions corresponding to the points in B. Under these conditions, IB-optimal choice behavior (as specified by *p*(*r*|*x*), the choice probabilities) is equivalent to adding softmax (logistic) choice noise to the exact posterior probability, *p*(*y*|*x*), such that higher values of *β* (proportional to *β*^∗^) correspond to less noise. This result can be equivalently formulated as the noisiness (imprecision) of an internal estimate of *p*(*y*|*x*) (inset, see text). **(D)** Certain heuristics (simplified inference strategies) can be understood as performing Bayesian inference on simplified encodings of the observations *X* and are characterized by their own “heuristic” IB bounds. **(E)** We can measure information capacity and predictiveness for individual decision-makers (points) and compare each to the full and heuristic IB bounds. We expect (cartoon data shown for illustrative purposes) that a substantial amount of individual variability (left) reflects variation in information capacity (here labeled “noise”) across the full and heuristic (i.e., biased) IB bounds (right; additional variability is shown as yellow points labeled “neither”).

Presumably, a decision-maker wants to maximize how closely their inferences match the true latent state. This match can be quantified as *I*(*R*; *Y*), the mutual information between *R* and *Y* (“inference predictiveness”). However, we assume it is costly to process information from *X* to form inferences. The amount of information used in this process can be quantified as *I*(*X*; *R*), the mutual information between *X* and *R* (“information capacity”). For the tasks we considered, limiting *I*(*X*; *R*) to reduce this cost (either voluntarily or due to inherent capacity limits) necessarily reduces the maximum-achievable *I*(*R*; *Y*). In other words, the less information is retained from the observations, the less information can be extracted about the latent state. This tradeoff is captured by the IB optimization problem:

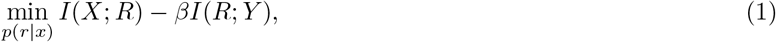

where *β* determines the relative importance of information capacity (minimized for smaller values of *β*) and inference predictiveness (maximized for larger values of *β*). The problem is solved by optimizing the choice probabilities *p*(*r*|*x*) [13]. Solving equation (1) over a range of *β* values (≥ 0) provides the IB bound, which defines the maximum possible *I*(*R*; *Y*) for a given *I*(*X*; *R*) (Fig. 2B). The exact nature of these relationships and the overall shape of the IB bound is unique for each task (as specified by the joint distribution *p*(*x, y*)).

Because we have full knowledge of *X*, *R*, and *Y* for each participant, we can estimate *I*(*X*; *R*) and *I*(*R*; *Y*) directly. By comparing these estimates to the IB bound, we can determine whether a given participant achieved the maximum possible performance given the information capacity they used. Individuals who lie on or near the bound optimally trade off performance with information-capacity limitations, which we term “information efficient”. Crucially, we can perform this analysis without requiring any knowledge about the strategies that individuals are using.

### 2.2 Capacity-limited, information-efficient inference implies evidence-based choice noise

A notable feature of the IB framework as it relates to human inference is its ability to distinguish noisy (information-efficient) inference from all forms of biased (information-inefficient) inference (Fig. 2B, E). Consider the choice probabilities *p*(*r*|*x*) that satisfy the IB optimization problem in general [13]:

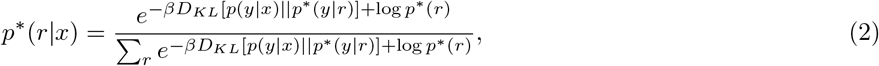

where *D_KL_* is the KL divergence. Because this expression is recursive (*p*^∗^(*y*|*r*) and *p*^∗^(*r*) depend on *p*^∗^(*r*|*x*)), the choice probabilities are computed numerically using an iterative algorithm (see Methods).

Equation (2) can be simplified to a familiar and more interpretable expression for many common cases (including our tasks; see Figs. B14, B16 for validation). Namely, when the IB-optimal decision-maker is equally accurate in inferring each latent state and makes all types of errors with the same probability (“accuracy symmetry”), and when the prior distribution over latent states *p*(*y*) is uniform (see Math Appendix), the probability of making the inference *r* takes the form of the standard softmax function:

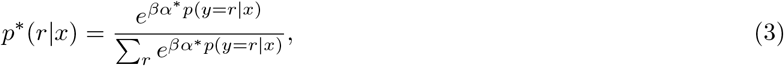

in which

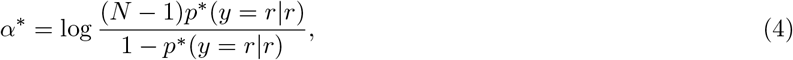

where *p*^∗^(*y* = *r*|*x*) is the posterior probability of the latent state corresponding to *r*, *p*^∗^(*y* = *r*|*r*) is the overall probability of making a correct inference (i.e., the accuracy), and *N* is the number of unique latent states. Because *α*^∗^ increases monotonically as *β* increases (see Math Appendix), each value of *β* (i.e., each point along the IB bound) corresponds to a unique value of *βα*^∗^, which we denote *β*^∗^. In short, (3) states that capacity-limited, information-efficient inference has effects that are equivalent to corrupting exact Bayesian inference with softmax (logistic) choice noise (blue-green points that fall along the IB bound in Fig. 2B, with corresponding psychometric curves in Fig. 2C). In other words, optimally reducing the information capacity of inference necessarily results in this classic form of evidence-dependent choice noise, with decreasing capacity corresponding to a greater noise magnitude (lower *β*^∗^).

Although this result tells us that any strategy for our tasks that is equivalent to Bayesian optimal inference corrupted by logistic choice noise falls on the IB bound, it does not prescribe a specific algorithm by which inferences should be formed. In principle, these algorithms can take many forms. One set of possibilities can be derived by assuming that the brain: 1) forms independent, potentially noisy estimates of the posterior probabilities of each latent state on a given trial; and 2) always chooses whichever latent state corresponds to the highest estimate. Under these assumptions, capacity-limited, information-efficient decision-makers behave as though they form noisy estimates *q_i_* of the posterior probabilities of each latent state *i* on each trial as follows:

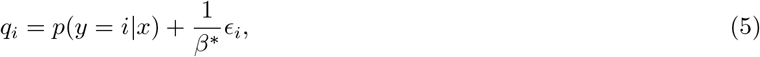

where the noise *ɛ_i_* is sampled independently from the standard Gumbel distribution for each *q_i_*. From this set of noisy estimates, the agent chooses the latent state with the highest probability estimate:

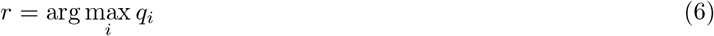

As established in prior work [20–22], such a process produces softmax-distributed choice probabilities (i.e., equation (3); Supplemental Fig. A1). As *β*^∗^ increases, each probability estimate *q_i_* becomes less noisy and converges to the true value of *p*(*y* = *i*|*x*), corresponding to higher information capacity and accuracy (Fig. 2C, inset). In the context of bounded rationality, people may be limited in their capacity or willingness to quench this noise, thus effectively lowering the precision of inference.

Although equation (3) describes the only way to be fully information efficient when *p*(*y*) is uniform and accuracy symmetry holds, other forms of evidence-dependent choice noise can be nearly as efficient. Commonly occurring cases include include posterior probabilities plus Gaussian noise, log posterior odds plus logistic noise [23, 24], and log posterior odds plus Gaussian noise [25]. IB curves constructed for these forms of noise nearly match the true IB bound for the tasks we considered (Supplemental Fig. A2).

Strategies that are less information efficient (i.e., fall further off the IB bound) are equivalent to incorrectly computing *p*(*y*|*x*). These biased strategies involve systematic errors in one’s beliefs about the statistical relationship between the observations and the latent state, which can include: 1) choosing one alternative (latent state) more often than the other, even though both have the same prior probability (Supplemental Fig. A3); 2) misestimating key variables (e.g., transition probabilities); or 3) making incorrect assumptions about the generative structure (e.g., believing there are statistical relationships like sequential dependencies when there are none).

An important form of the last are strategies based on simplifying assumptions about the generative structure, otherwise known as heuristics [9, 26] (Fig. 2D). Certain heuristics are equivalent to inferring the latent state using a “heuristic encoding” of the observations. In contrast to the “full encoding” (*X* in its original form), a heuristic encoding might, for example, include only the observations providing the most information about the latent state. Although this is a form of compression, it is not necessarily information efficient. For our tasks, equation (3) implies that the optimal compression of *X* incorporates all parts of *X*, weighted according to their relative informativeness. Heuristic encodings that ignore or improperly weigh informative parts of *X* are thus not IB-optimal compressions of *X*.

Nevertheless, heuristic strategies can also follow principles of bounded rationality evident as an information bottleneck. If a heuristic is equivalent to computing *p*(*y*|*x*) using a heuristic encoding of *X*, then a “heuristic IB bound” can be constructed by using the heuristic *X* in place of the full *X* (Fig. 2E). Thus, decision-makers on such a heuristic IB bound are performing information-efficient inference much like those along the “full” IB bound, albeit with the simplified heuristic encoding of the observations. Like for the full IB bounds, solutions to the IB problem for the heuristics of interest for both tasks satisfy the conditions necessary for equation (3) to hold (Figs. B15, B17).

### 2.3 Behavioral variability across individuals reflects information-efficient inference

For the bead-prediction task, performance varied considerably across participants but tended to be information efficient, given their measured information capacity and apparent strategy. Specifically, inference predictiveness varied from near zero, corresponding to random guessing or choosing only one option, to nearly perfect, matching ideal-observer levels of 0.66 bits of predictiveness or 93.9% correct (y-axis in Fig. 3A). Despite this variability, 56 out of 64 participants had choice behavior that fell on (i.e., had inference predictiveness with 95% bootstrap confidence intervals that overlapped) either: 1) the full bound, which characterizes how maximum possible performance changes as a function of information capacity (equivalent to varying the amount of logistic choice noise) using fully optimal inference (45 out of 56, blue-green points in Fig. 3A, psychometric functions in the left column of 3B); or 2) the heuristic “one-back” bound, which characterizes how maximum possible performance changes as a function of information capacity using a specific heuristic that involved using only the most recent bead draw (11 out of 56, red points in Fig. 3A, psychometric functions in the right column of 3B; participants that overlapped with neither are shown in yellow).

**Fig. 3:**
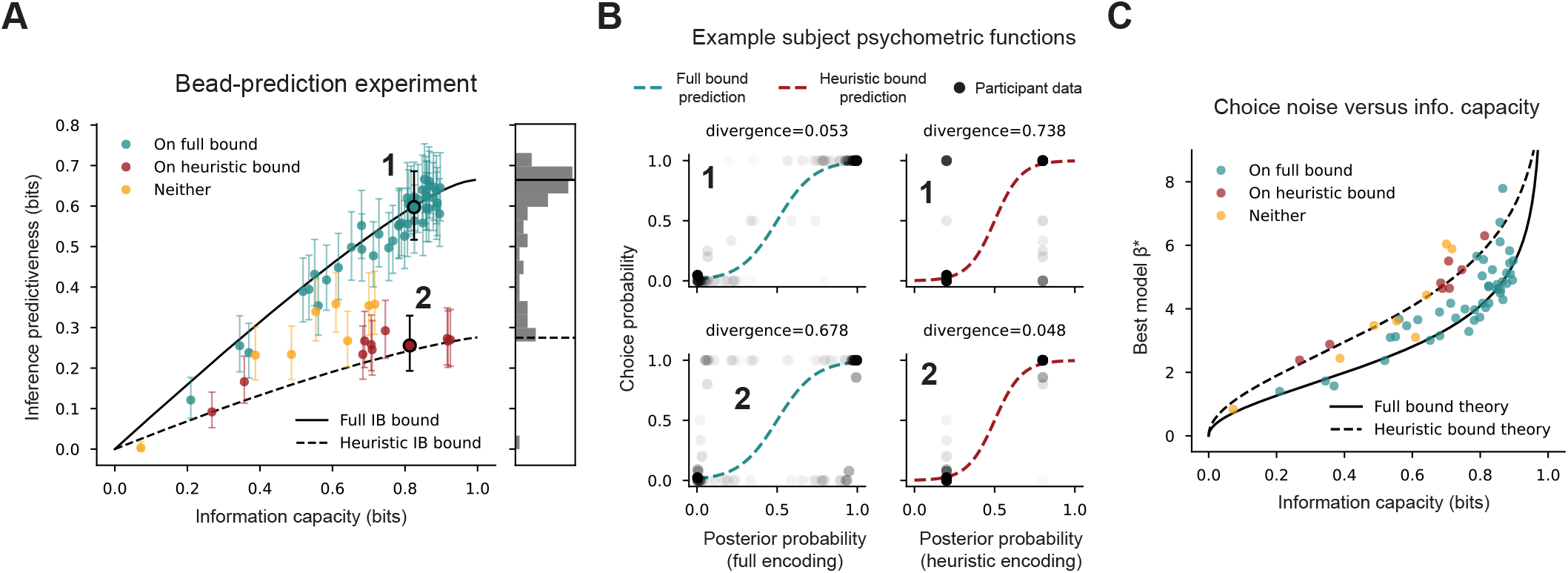
Human behavior on the bead-prediction task. **(A)** Individual participants (points, N=64) plotted against the full and heuristic IB bounds for block 2 of the bead-prediction task. Participants were classified based on the closest bound to which their 95% confidence interval (error bars) on predictiveness overlapped (or neither; see legend). The histogram to the right gives the distribution of information efficiency (predictiveness normalized by the value of the full IB bound at a given participant’s information capacity). A substantial amount of individual variability can be explained by variation in information capacity along either the full or heuristic IB bound. **(B)** Example psychometric functions for the two participants highlighted in A. In each subplot, each shaded black point gives the probability the participant chose jar 1 across all trials with the same sequence of 7 bead draws as a function of the posterior probability (*p*(*y*|*x*)) for that particular sequence. Shading indicates the relative frequency of that particular bead sequence. The posterior was computed using either the full encoding of the observations *X* (most recent 7 bead draws, left panels) or the one-back heuristic encoding (only the most recent bead drawn, right panels). Dotted curves are the IB-optimal psychometric function, computed using the given enocoding and the participant’s information capacity. The KL divergence above each panel indicates the mismatch between the participant’s choice probabilities and the IB-predicted choice probabilities associated with the given psychometric function (participant 1 shows a better match to the full-bound prediction, participant 2 shows a better match to the heuristic-bound prediction, consistent with their positions in IB space in A). **(C)** Logistic noise magnitude of the best-fitting model (*β*^∗^) plotted versus the information capacity for each participant (points) compared to theoretical relationships (lines, as indicated). Choice noise is closely related to information capacity as predicted by equation (3) in a bound-specific manner.

As expected, participants along the full bound exhibited choice probabilities that matched the full encoding posterior probabilities plus logistic noise (example participant 1, Fig. 3B), whereas participants along the one-back bound more strongly matched the one-back encoding posterior probabilities plus logistic noise (example participant 2, Fig. 3B). The degree of evidence-dependent choice noise was closely related to information capacity (equation (3)) for participants’ respective bounds (Fig. 3B,C). This relationship converged to the precise theoretical prediction as individual choice behavior more strongly matched either the full or one-back bounds (Supplemental Fig. A4).

We compared these results to fits to the data of the fully optimal model (Bayesian inference using the 7 most recent bead draws) and one-back model (one free parameter each; choice noise via *β*^∗^). In general, participant behavior was more consistent with the fully optimal model along the full IB bound and with the one-back heuristic along the one-back bound (Fig. 4A). We also performed a more extensive model comparison that included a “subjective-hazard model.” This model used the preceding sequence of bead draws like the fully optimal strategy but made the hazard rate of jar switches an additional free parameter. Thus, the subjective-hazard model accounted for the possibility that participants attempted to implement the fully optimal strategy but misestimated the hazard rate relative to the true value of 0.01 (a form of non-heuristic bias). We ruled out another form of non-heuristic bias, choosing one latent state more often than the other on average (see Supplemental Figs. A5, A6, A7, A8).

**Fig. 4:**
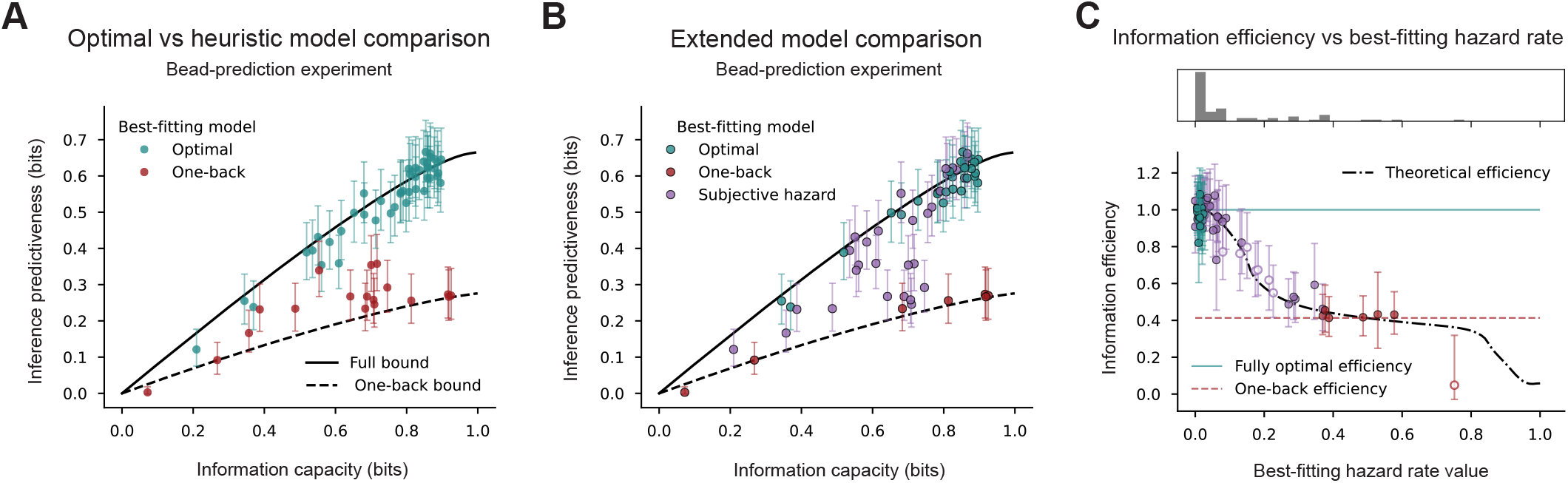
Model comparisons, bead-prediction task. **(A)** IB plots of human behavioral data for the bead-prediction experiment, with participants colored by best-fitting model in a comparison between the fully optimal model (Bayesian inference using the 7 most recent bead draws) and the one-back model (Bayesian inference using only the most recent draw). **(B)** Same as A, but including the “subjective-hazard” model (like fully optimal, but letting the hazard rate be a free parameter). **(C)** Information efficiency (predictiveness divided by the value of the full IB bound at a given information capacity, shown as a histogram above) plotted versus best-fitting hazard rate from the subjective-hazard model for each participant. Unfilled points denote participants that were not on either the full or heuristic IB bound. The black dash-dotted line gives the theoretical efficiency (computed at the information capacity of each participant and then averaged across participants) of the subjective-hazard model for a given hazard rate. The blue-green and red-dashed lines give the information efficiencies of the fully optimal and one-back strategies, respectively (averaged across participants for one-back). Of the participants best fit by the subjective-hazard model, most had fitted hazard rate values that fell within a relatively information-efficient range.

For this extended model comparison, many participants along the full and one-back bounds were best fit by the corresponding model (26 out of 45 on the full bound, 6 out of 11 on the one-back bound; Fig. 4B). The remainder near the full IB bound were best fit by the subjective hazard model (19 out of 45). These participants appear to have used misestimated hazard-rate values that nonetheless supported information-efficient inference. Consistent with this idea, many participants best fit by the subjective-hazard model (purple points, Fig. 4C) tended to be concentrated in the region of parameter space (particular hazard rate values) that was highly information efficient (histogram, 4C). Only seven participants fell in between the two bounds (i.e., did not have confidence intervals that overlapped with either one). Their behavioral data were best fit by the subjective-hazard model (Figure 4B), with hazard-rate parameter values falling within relatively inefficient ranges from this perspective (unfilled points, Figure 4C).

To examine the generality of these trends and their dependence on the particular task statistics, we tested participants using three versions of the horse-prediction task. These versions differed in terms of the relative amount of evidence (quantified by log-likelihood ratios, see Methods) provided by the “weak” and “strong” shapes (the number of weak shapes required to equal the evidence strength of a single strong shape, which we term the W-S ratio). One had a “low” W-S ratio (1.3:1), which implies that the weak and strong shapes provided nearly the same amount of evidence. Accordingly, a heuristic that equally weighs the strong and weak shapes (“equal-weights heuristic”, which assumed a W-S ratio of 1:1) is nearly equivalent to the fully optimal strategy in terms of information-efficiency. The remaining two had either an “intermediate” (2.5:1) or “high” (6.3:1) W-S ratio, for which the equal-weights heuristic would be increasingly less information-efficient.

Across all versions of the horse-prediction task, individual differences in behavior were consistent with substantial variation in information capacity that fell along either the full IB bound or the equal-weights heuristic IB bound (Fig. 5A,D,G). Moreover, this variation in information capacity was closely related to evidence-dependent choice noise, as predicted by equation (3) (Fig. 5B,C,E,F,H,I). However, the position of the heuristic bound, and thus the distribution of participants in IB space, varied systematically by task condition. Specifically, for the low W-S ratio version, nearly all participants (29 out of 30) had choice behavior that fell along the full (and nearby equal-weights heuristic) bound (Fig. 5A). As expected, psychometric functions were similar when computed with respect to the fully optimal or equal-weights heuristic posterior probabilities (Fig. 5B). In contrast, for the high W-S ratio version, fewer participants (16 out of 30) had choice behavior that fell near the full bound, with most of the remaining participants (11) near the much lower equal-weights heuristic bound (Fig. 5G). Under these conditions, the choice behavior of participants along the two different bounds were highly dissimilar in terms of their psychometric functions (Fig. 5H, 1 versus 2). For the intermediate W-S ratio version, performance was intermediate to these trends (Fig. 5D-F).

**Fig. 5:**
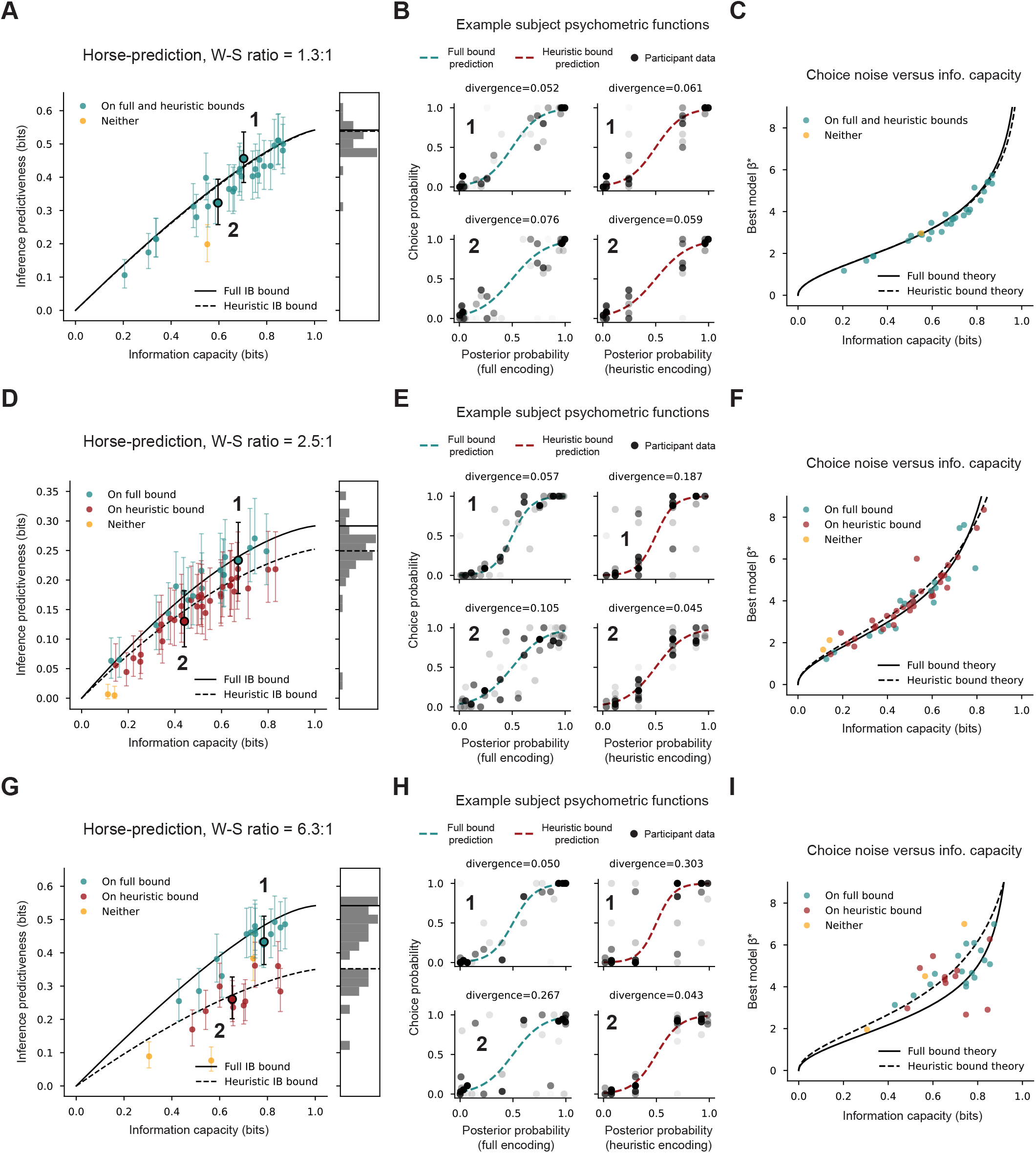
Human behavior on the horse-prediction task. We used three versions of the horse-prediction task that differed in terms of the “W-S ratio,” which gives the number of weak shapes required to equal the evidence strength of a single strong shape. Rows use the same conventions as in Figure 3. In the psychometric function plots, each point corresponds to a unique combination of shape cards (the full encoding of *X* in this task). **(Top row)** Low W-S ratio (1.3:1) (N=30). The equal-weights heuristic is nearly indistinguishable from optimal inference, and thus both correspond to similar IB bounds (A and C) and psychometric functions (B). **(Middle row)** Same as top row for intermediate W-S ratio (2.5:1) (N=53). The equal-weights heuristic is less information efficient than optimal inference but still relatively close to the full IB bound. **(Bottom row)** High W-S ratio (6.3:1) (N=30). The equal-weights heuristic is substantially less information efficient (the heuristic bound is lower). Despite the large change in task statistics across experiments, individual differences largely reflected variability in information capacity along either the full or equal-weights IB bound. Furthermore, choice noise was closely related to information capacity as predicted by equation 3 across all three experiments.

Like for the bead-prediction task, a model-fitting analysis further supported the idea that, for the given strategy, information capacity, and task condition, people tended to be information efficient. Specifically, for all versions of the horse-prediction task, behavior was most consistent with the fully optimal model along the full IB bound and with the equal-weights heuristic along the heuristic bound, when these bounds were distinct (Fig. 6A). When we included additional models in the model comparison (the “ignore-weak heuristic” computed the evidence from the strong shapes only; the “subjective-ratio model” made the W-S ratio a free parameter and thus served to capture non-heuristic bias as in the bead-prediction task), participants along the equal-weights bound were predominantly best-fit by the equal-weights model across all versions with distinct bounds (Fig. 6B). Meanwhile, participants along the full bound but best fit by the subjective-ratio model again tended to concentrate in regions of parameter space (particular W-S ratios) that were highly information efficient (Fig. 6B,C). Crucially, such participants found this efficient region in each version despite the fact that its location changed in a version-dependent manner (Fig. 6C).

**Fig. 6:**
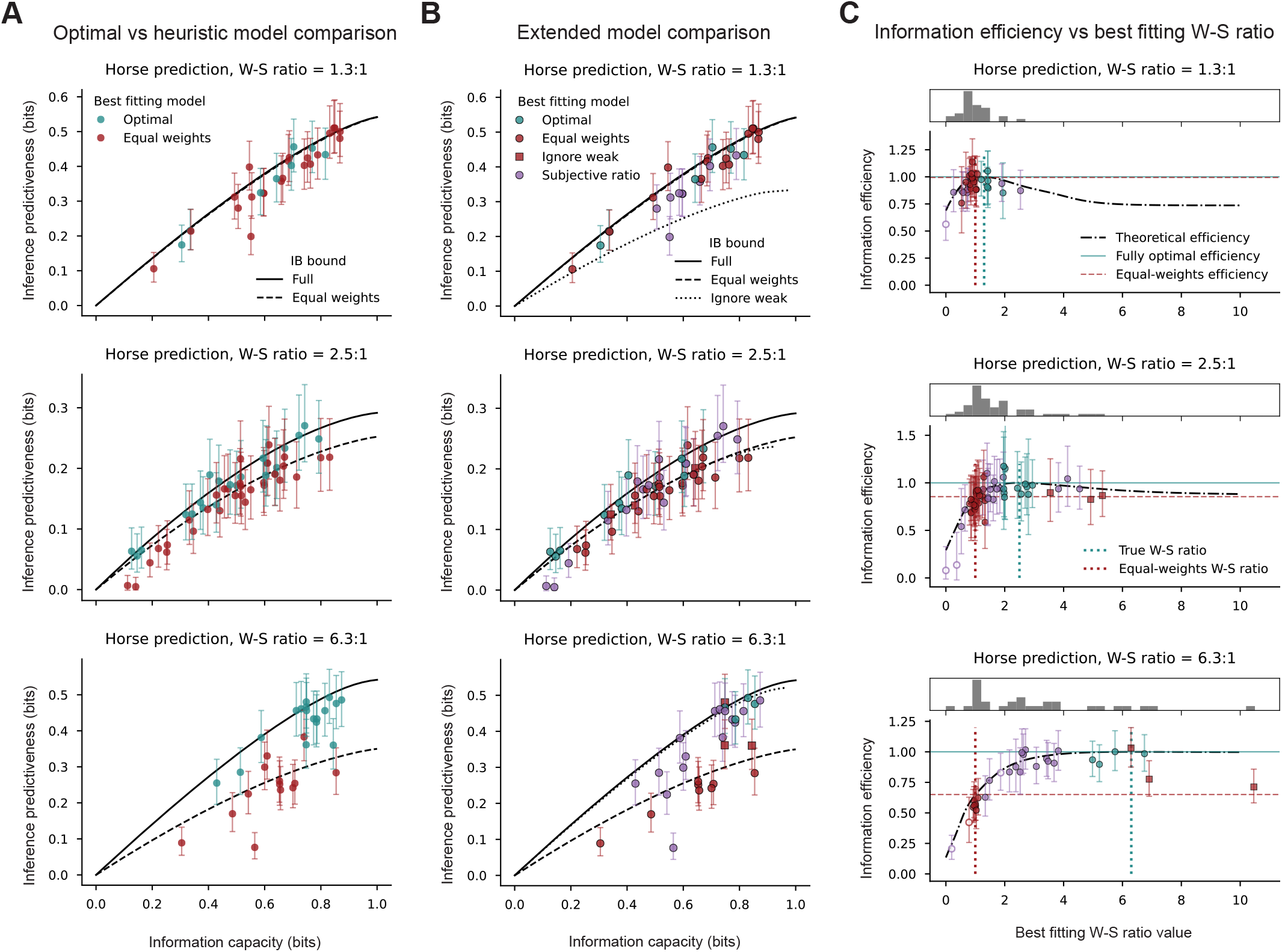
Model comparisons, horse-prediction task. **(A)** IB plots of human behavioral data for the three versions of the horse-prediction task (rows), with participants colored by best-fitting model in a comparison between the fully optimal and equal-weights models. **(B)** Same as A, but for an extended model comparison including the “ignore-weak” (a heuristic ignoring the weak shapes) and “subjective-ratio” (like fully optimal, but letting W-S ratio be a free parameter) models. Like for the bead-prediction experiment, many participants were best fit by the model corresponding to the bound they were on. Many also appeared to be using optimal inference but with a misestimated W-S ratio. **(C)** Information efficiency (predictiveness divided by the value of the full IB bound at a given information capacity, shown as a histogram above) plotted versus best fitting W-S ratio from the subjective-ratio model for each participant. The black dash-dotted line gives the theoretical efficiency (computed at the information capacity of each participant and then averaged across participants) of the subjective-ratio model for a given W-S ratio. The blue-green and red-dashed lines give the information efficiencies of the fully optimal and equal-weights strategies respectively (averaged across participants for equal-weights). Although many participants were better fit by the subjective-ratio model, most fell within relatively information-efficient regions of parameter (W-S ratio) space. This phenomenon was especially apparent in the high W-S ratio experiment, for which a broad range of ratio estimates supported information-efficient inference.

These model comparisons also showed systematic differences in the tendency to use particular heuristics for the three versions of the task. In particular, the proportion of participants best fit by the equal-weights model decreased progressively (from 50% to 41.5% to 26.6%) as the W-S ratio increased and this strategy became increasingly inefficient. Conversely, the proportion of participants best fit by the ignore-weak model increased progressively (from 0% to 5.7% to 10%) as the W-S ratio increased and this strategy became increasingly efficient. Thus, individual variability reflected not only differences in information capacity for a given strategy, but also differences in strategy use across different task versions to maintain information efficiency.

### 2.4 Behavioral adjustments by individuals maintain information efficiency

Having shown that principles of information efficiency govern variability in performance across participants, we then tested if and how these same principles relate to behavioral adjustments made by individual participants in response to different task conditions. We tested three conditions that commonly affect performance accuracy: learning, speed-accuracy trade-offs, and reward incentives (Fig. 7). As detailed below, we used the IB to show that changes in accuracy (measured as inference predictiveness) were accompanied by associated changes in information capacity that tended to maintain information efficiency (i.e., that changed both capacity and predictiveness in a way that moved along the IB bound, as opposed to, for example, improving predictiveness while maintaining similar information capacity).

**Fig. 7:**
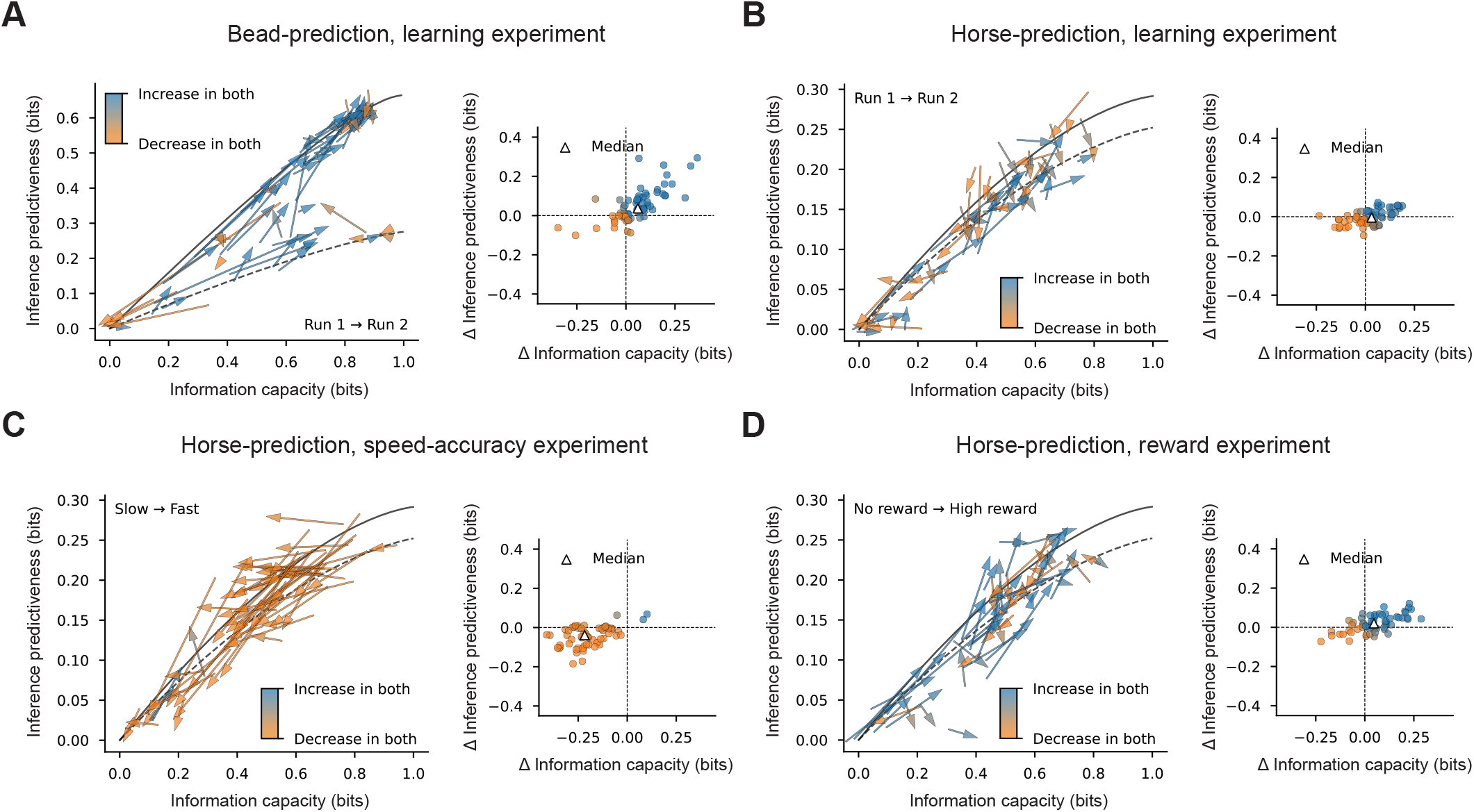
Behavioral-adjustment experiments. **(A)** Left, IB plot for the bead-prediction learning experiment showing the change in information capacity and inference predictiveness from run 1 (arrow tail) to the identical run 2 (arrow head) for each participant. Colors indicate if both increased (blue), both decreased (orange), or neither (gray) occurred. Right, Change in inference predictiveness versus change in information capacity plotted for each participant (points). **(B)** Same as A, but for the horse-prediction learning experiment. **(C)** Same as A, but for the horse-prediction speed-accuracy experiment. Arrows point from the slow decision-time condition (arrow tails) to the fast decision-time condition (arrow heads). **(D)** Same as A, but for the horse-prediction reward experiment. Arrows point from the no-reward condition (arrow tails) to the high-reward condition (arrow heads). In all four cases, participants typically changed capacity and predictiveness in a manner that maintained information efficiency (stayed on an IB bound), with an overall tendency for both to increase (A and D), decrease (C), or neither (B).

Repeated performance resulted in different patterns of behavior for the two tasks, but in both cases reflected adjustments that maintained (or even increased) information efficiency. For the bead-prediction task, there were substantial learning effects from run 1 to run 2, corresponding to increases in inference predictiveness for most participants (47 out of 68 increased, median change = 0.037 bits; two-sided Wilcoxon signed-rank test: *n* = 68, *T* = 455.0, *p* = 1.9 × 10^−5^). These changes were accompanied by increases in information capacity (48 out of 68 increased, median change = 0.060 bits; *n* = 68, *T* = 526.0, *p* = 7.7 × 10^−5^) that in most cases corresponded to movement along the associated bound in IB space (i.e., increases in capacity when predictiveness increased and decreases in capacity when predictiveness decreased; Fig. 7A). In addition, several participants moved from near the one-back heuristic bound upward toward the full IB bound, resulting in a slight overall increase in information efficiency (predictiveness divided by the value of the full IB bound at a given information capacity) from run 1 to run 2 (38 out of 63 increased, median change = 0.014; *n* = 63, *T* = 658.0, *p* = 0.017).

For the horse-prediction task, changes in performance were more variable from run 1 to run 2, including increases and decreases in both information capacity (36 out of 60 increased, median change = 0.032 bits; *n* = 60, *T* = 663.0, *p* = 0.064) and inference predictiveness (27 out of 60 increased, median change = −0.0045 bits; *n* = 60, *T* = 840.0, *p* = 0.58) across participants (Fig. 7B). This difference between tasks likely reflects the more extensive training we gave participants on the horse-prediction task before testing, thus limiting systematic improvements with learning on the learning blocks (group mean accuracy = 70.0% on the training block, 70.4%, on learning block 1, and 70.3% on learning block 2). Nevertheless, individual behavioral changes across runs for the horse-prediction task were associated with adjustments along either the full or heuristic IB bound. That is, information capacity and inference predictiveness increased or decreased concurrently in a manner that maintained a given participant’s information efficiency for their given, relatively stable, strategy across the two runs (23 out of 53 increased, median change = −0.025; *n* = 53, *T* = 558.0, *p* = 0.16).

Restricting decision times on the horse-prediction task (randomly interleaving trials with 0.75 s or 1.75 s viewing times) resulted in systematic reductions in accuracy, consistent with a speed-accuracy trade-off, that also maintained information efficiency (Fig. 7C). Specifically, restricting decision time substantially reduced both the measured information capacity and inference predictiveness for nearly every participant (information capacity: 58 out of 60 decreased, median change = −0.22 bits; *n* = 60, *T* = 19.0, *p* = 4.2 × 10^−11^; inference predictiveness: 48 out of 60 decreased, median change = −0.039 bits; *n* = 60, *T* = 194.0, *p* = 1.1 × 10^−7^). As with learning, changes in these measures reflected adjustments along the IB bound, such that participants tended to maintain their information efficiency across the two conditions (31 out of 57 increased, median change = 0.090; *n* = 57, *T* = 628.0, *p* = 0.11). Thus, decreases in performance caused by limiting decision time reflected decreased information capacity rather than switches to simpler, but less information-efficient, strategies.

Likewise, changes in incentives caused changes in behavior that maintained information efficiency on the horse-prediction task (Fig. 7D). Participants completed one block with no rewards (positive or negative) for their choices and one block with double the reward amounts used in all of the other horse-prediction experiments (block order was randomized between participants). The two blocks were otherwise identical, including the observations shown on each trial. The participants tended to increase both information capacity and inference predictiveness when given reward incentives (two-sided Wilcoxon signed-rank test; information capacity: 43 out of 60 increased, median change = 0.047 bits; *n* = 60, *T* = 448.0, *p* = 5.9 × 10−4; inference predictiveness: 40 out of 60 increased, median change = 0.020 bits; *n* = 60, *T* = 486.0, *p* = 0.0016; Linear mixed-effects model; information capacity: *n* = 60, *β* = 0.058, *t* = 3.7, *p* = 2.6 × 10−4; inference predictiveness: *n* = 60, *β* = 0.021, *t* = 3.5, *p* = 4.3 × 10^−4^) in a manner that maintained information efficiency (32 out of 55 increased, median change = 0.019; Wilcoxon signed-rank test; *n* = 55, *T* = 619.0, *p* = 0.21; Linear mixed-effects model; *n* = 55, *β* = 0.025, *t* = 0.86, *p* = 0.39).

## 3 Discussion

Human choice behavior is characterized by substantial variability between and within individuals. Here, we showed that this variability can be structured in principled ways that reflect limited-capacity, but information-efficient, inference. We used the IB framework to show that information efficiency is maximized on the “full” IB bound, which can be effectively described as fully optimal inference corrupted by a certain form of choice noise that governs the relationship between information capacity and inference predictiveness. Similar relationships can be derived for other, lower bounds that represent information-efficient implementations of suboptimal (heuristic) strategies. We found that behavioral variability across individuals for particular tasks, and across task conditions within individuals, tended to be distributed along these full and heuristic IB bounds. Together, these results imply that human inference not only efficiently maximizes performance given information capacity, but also can flexibly adjust this efficient capacity-accuracy tradeoff to alter the balance between performance and resource frugality.

For the inference problems we considered, information-efficient decision-makers (those on an IB bound) behaved as if they were computing the exact posterior probability (using the full encoding of the observations *X* or a simplified one corresponding to a heuristic) and then corrupting their choices with an amount of logistic noise determined by their information capacity. This form of evidence-dependent choice noise is commonly observed in decision-making tasks and has been interpreted in terms of inherent neural noise [22, 25, 27], natural tendencies toward exploratory behaviors [28], and other factors. Our results imply that this noise might also reflect a form of bounded rationality that balances inherent tradeoffs between performance and reducing the cost of information capacity. Our results also show that this noise can be adjusted by individuals while maintaining efficiency under a variety of conditions, including changes in incentives that imply at least some volitional control over these adjustments.

Our demonstration that IB-optimal choice behavior is equivalent to performing exact Bayesian inference and then adding noise during the process of making a choice does not imply that this is the mechanism by which humans form capacity-limited inferences. In fact, this mechanism would generally be highly resource inefficient and therefore seems unlikely. In contrast, there are other mechanisms that would produce equivalent behaviors (i.e., result in the same, IB-optimal choice probabilities) but use resources more efficiently. For example, the additive noise model (equations (5) and (6); [20–22]) suggests that reduced information capacity arises from imprecision (noise) in the computation of posterior probabilities. This idea is consistent with previous suggestions that suboptimal human decision-making reflects limits on the precision of inference [25, 29]. We extended these findings by demonstrating that inference imprecision can reflect general limitations in the amount of costly computational resources the brain is able (or willing) to devote to a particular problem.

Our speed-accuracy findings provide more clues as to the nature of information-efficient, capacity-limited inference in the brain. Under the interpretation that information capacity indexes the precision of inference, our findings that information capacity declined sharply when decision time was restricted implies that this precision results from a process that unfolds over time. This idea is consistent with commonly used decision-making models that are based on the temporal accumulation of noisy evidence sampled sequentially over time (which can occur when the evidence itself is presented sequentially or, like in our task, presented all at once but then processed over time) [30]. Ending this process early results in a less precise representation of the integrated evidence and less accurate inference. We showed that this temporally dynamic process tends to be information efficient. That is, akin to using perfect integration to make the most effective use of noisy information that is presented sequentially over time, the brain appears to use a similarly efficient process to dynamically sample and use information that is presented all at once to make inferences. This idea also suggests that increasing information capacity may be costly in terms of time, a resource used for many decision-making problems [7, 31].

Other studies have used related frameworks to explore relationships between information capacity and performance under a variety of conditions. One line of work [32–36] investigated these tradeoffs for behavioral policies that aim to maximize reward (as opposed to state inferences to maximize accuracy in our study). Under those conditions, human decision-makers vary considerably in information capacity (“policy complexity”) but often lie near the capacity-dependent, reward-maximizing bound, similar to our findings. Moreover, reducing information capacity is equivalent to increasing the magnitude of logistic (softmax) choice noise added to the state-action (*Q*) values when the marginal distribution of optimal actions is uniform. When this marginal is not uniform, optimal reductions in information capacity correspond to not just increasing choice noise, but also biasing the marginal choice distribution toward the most rewarding choice, on average (the IB framework makes the same prediction for our applications; see Supplemental Figs. A9, A10).

An analogous set of findings has been made in the rational-inattention literature in economics [37–40]. This line of work considers decision-makers who aim to maximize the utility of their choices but are limited in how much information they can process to infer the optimal choice. Under this framework, optimally reducing information capacity is again associated with increasing logistic choice noise and bias towards the most rewarding choices (when the marginal distribution of optimal actions is not uniform). Although empirical tests of this framework have not compared individuals to an optimal bound, numerous studies have found that the rational-inattention model can account for certain properties of human choice behavior for a variety of problems [40–43]. Thus, for our problem, and other similar one, optimal changes in information capacity correspond to varying the amount of evidence/reward-based choice noise and potentially bias toward certain choices.

These findings contrast notably with the nature of optimal information compression in other problems as revealed by the IB framework. Namely, movement along an IB bound sometimes corresponds to varying the coarseness of approximations, analogous to how digital images can be compressed to varying degrees of “pixelation”. For example, when the observed signal (*X*) and the inference target (*Y*) are multivariate Gaussian variables, optimally decreasing the information capacity essentially corresponds to using fewer dimensions to store information in the compressed signal (*R*) [44]. In our context, movement along the IB bound does not correspond to a similar notion of degree of approximation. Instead, we find that one form of approximate inference, heuristics, corresponds to distinct, lower IB bounds using simplified versions of *X*. This discrepancy hints at the kinds of specializations that the brain may use to implement compression in a way that maintains information-efficient inference.

Our finding that many human decision-makers fall on heuristic bounds implies that the IB principle and the form of bounded rationality it represents cannot explain individual variability on its own. In addition, there are likely costs associated with strategies that fall on the full IB bound. One potential source of these costs is related to the computational complexity of those strategies. This idea is central to many studies of bounded rationality in decision-making, which include the finding that human performance relative to optimal declines as the computational complexity of particular decision-making problems increases [45, 46]. Accordingly, people often rely on heuristic strategies, especially as problems become more complex [8, 9]. The use of notably information-inefficient, simplified inference strategies in our data likely reflects the same form of bounded rationality. More work is needed to understand how costs related to information capacity, computational complexity, and other factors together serve to constrain human decision-making.

In summary, our work identifies fundamental principles of human inference that structure a substantial amount of individual variability in human choice behavior. We showed that human inference can be largely information efficient, maximizing performance given information capacity for a particular observation encoding scheme (full or simplified). Moreover, the information capacity of human inference can be flexibly adjusted while maintaining information efficiency, enabling adaptive switching between high-performance, high-cost and cost-saving, but less-accurate, inferences. These findings help to establish information use as a fundamental driver of the bounded rationality that governs our ability to process information and interact with the world.

## 4 Methods

### 4.1 Human psychophysics

Human subject protocols were approved and determined to be exempt by the University of Pennsylvania Internal Review Board (IRB protocol 844474). Participants provided consent online before they began the session.

All experiments were performed on the online platform Prolific. We used the following inclusion criteria to recruit participants: 18–65 years old (self reported), fluent in English (self reported), be of US nationality (self reported), have a Prolific task approval rate of 99–100%, and must have completed at least 20 tasks on Prolific. During data collection, we excluded any participant who did not complete the entire experiment. We then collected additional data until we reached our predetermined number of participants (see below). For the horse-prediction experiments, we additionally identified and excluded from additional analyses any participant who performed too close to chance, defined as an information capacity of *<*0.05 bits in all experimental conditions. We collected data from additional participants until we reached the desired number who passed this performance criterion.

#### 4.1.1 Horse-prediction task

The horse-prediction task (a variant of the weather-prediction task [4, 19]) required inferences about a latent state (which of two horses would win the race) based on multiple independently sampled observations with varying evidence strengths (likelihood values). The winning horse on each trial was chosen at random (50% probability for each). Based on which horse would win on a given trial, five shapes were presented simultaneously to the participant. These shapes were sampled independently with replacement from one of two probability distributions over four unique shapes.

We parameterized these likelihood distributions by specifying the log-likelihood ratios (LLRs; ratio between the likelihood given one horse versus the likelihood given the other horse; base 10) of the four shapes in the form of [-(W-S ratio)×weak, -weak, weak, (W-S ratio)×weak], where “weak” gives the LLR of the weak shapes and “W-S ratio” gives the ratio between the LLR of a strong shape and the LLR of its corresponding weak shape. To specify a unique pair of likelihood distributions, we were required to set a reference probability that we refer to as *p*_1_, the likelihood corresponding to the first shape in our list of LLRs, -(W-S ratio)×weak. Given these specifications, two shapes represent strong and weak evidence that one horse will win, and the other two represent strong and weak evidence that the other will win. The overall strength of the evidence for and against each latent state on a given trial is specified by the sum of the LLRs of the five shapes that appear (shown in five fixed positions equidistant from the center of the screen and from each other).

At the beginning of each session, participants were told which shapes provided evidence that either the green or purple horse would win. Participants were also told which shapes were strongly indicative of their horse and which were weakly indicative, although the exact likelihood values were not specified. Finally, participants were informed that sometimes the horse not favored by the shapes would win. The statistical relationships between the specific horses and shapes were randomized for each participant, along with the order of trials. Each participant then completed 100 training trials in which they were given unlimited time to view the shapes and make their decisions, followed by 10 “demo” trials in which they were given limited time to view the shapes and limited time to make their decisions, as would be the case in the testing trials. Finally, they completed 700 “testing” trials with generative statistics that matched that of the demo trials for the given task variant. The first 100 trials from this block were discarded to minimize learning effects within a block.

The green and purple horses each won in exactly one half of the testing trials. The set of trials comprising each half, as specified by the combination of five shapes that appeared, were chosen to match the relative frequencies of occurrence of each shape combination under each horse. For example, if a shape combination had an 8% probability of being generated given a particular horse winning, there were 12 trials with that shape combination within a set of 150 trials. Specifically, we multiplied the probability of a trial type (shape combination) by the number of trials, then rounded this number. We summed these rounded numbers together across all trials. If this sum was over the desired number of trials, took one trial away from each trial type in order from most common to least until the desired number was reached. If the sum was under, we added one trial to each trial type in order from least to most common (but only for trials ≥1 trial).

This method of construction was performed for each separate condition in an experiment for two reasons. First, it ensured that the set of trials was identical across participants and across conditions for the same participant, allowing us to compare directly measured values of information capacity and inference predictiveness (Supplemental Fig. A11). Second, with a relatively limited number of trials, it is possible to randomly generate a trial set that makes even information-efficient decision-makers simulated with equation (3) look information-inefficient (i.e. fall off of the IB bound) (Supplemental Fig. A11). Thus, we matched the frequencies of different shape combinations within the testing trials to the theoretical frequencies to minimize this distortion.

For the demo and testing trials, the shapes were shown for a limited time (that varied depending on experiment) before being masked. Once the shapes were masked, participants had 0.5 s to indicate which horse they believed would win. If a response was provided within this 0.5 s window, participants saw which horse actually won and thus whether they were correct. On trials in which the participant failed to provide a response in time, the trial ended and the participant was not shown the winning horse. These incomplete trials were added to the end of the block to be completed again. This process was repeated only three times before the block ended, and any remaining trials remained uncompleted (at least 96.3% of trials were completed by all participants in all conditions, with the exception of one participant who completed 85.3% of trials in the fast condition of the speed-accuracy experiment).

Participants earned a base payment of $9 for completing the experiment and could earn a bonus payment based on their level of performance on the testing trials. These bonuses were incremented by a particular amount for each correct response and decremented by the same amount for each incorrect response (but could not go below 0). The specific amount differed for different tasks. These amounts were added/subtracted from a running sum that started at $0 on each separate block of testing. The final sums for each separate block were added together to get the final bonus payment amount. We chose the bonus amount to span the range from random behavior (∼$0) to fully Bayes-optimal behavior (∼$5). All horse-prediction experiments were implemented using PsychoPy v2023.2.2 [47] and hosted on Pavlovia.

##### Base experiment, W-S ratio=1.3:1, small difference in weak vs. strong evidence

30 participants completed the task with the following parameters: weak=0.45, W-S ratio=1.3, *p*_1_=0.06, such that the LLRs of the four shapes were [−0.585, −0.45, 0.45, 0.585] (and thus the likelihood of drawing the first shape, corresponding to the LLR of −0.585, was p=0.06). Pre-testing consisted of a single set of 100 trials, followed immediately by a single block of 600 testing trials. Viewing time was 1.5 s. The bonus increment was ±$0.012/trial. Only one additional participant was collected to meet the performance criterion (information capacity ≥0.05 bits).

##### Base and learning experiments, W-S ratio=2.5:1, moderate difference in weak vs. strong evidence

60 participants completed the task with the following parameters: weak=0.2, W-S ratio=2.5, *p*_1_=0.08, such that the LLRs of the four shapes were [−0.5, −0.2, 0.2, 0.5]. Pre-testing consisted of a single set of 100 trials. Testing consisted of 600 trials, broken up into two sequential, identical blocks of 300 trials each. Given that there was no effect of learning (block 1 versus block 2) on information capacity and predictiveness, blocks 1 and 2 were combined and analyzed as part of the systematic evaluation of the effect of W-S ratio on participant behavior (“version with W-S ratio = 2.5:1”). In line with our other experiments, participants with an information capacity *<* 0.05 bits for the combined trial set were excluded from analyses (7 out of 60). Viewing time was 1.5 s. The bonus increment was ±$0.012/trial. Eight additional participants were collected to meet the minimum information capacity criterion.

##### Base experiment, W-S ratio=6.3:1, large difference in weak vs. strong evidence

30 participants completed the task with the following parameters: weak=0.18, W-S ratio=6.3, *p*_1_=0.02, such that the LLRs of the four shapes were [−1.134, −0.18, 0.18, 1.134]. Pre-testing consisted of a single set of 100 trials, followed immediately by a single block of 600 testing trials. Viewing time was 1.5 s. The bonus increment was ±$0.012/trial. Four additional participants were collected to meet the performance criterion.

##### Speed-accuracy experiment

60 participants completed the task with the following parameters: weak=0.2, W-S ratio=2.5, *p*_1_=0.08. Testing consisted of 600 trials, with 300 trials of the “slow” condition (1.75 s viewing time) and 300 trials of the “fast” condition (0.75 s viewing time). In both cases, participants had 0.5 s to respond once the shapes were masked. Slow and fast trials were interleaved randomly within the 600-trial block. Pre-testing consisted of 50 slow and 50 fast trials, all interleaved randomly. The bonus increment was ±$0.012/trial. Nine additional participants were collected to meet the minimum information capacity criterion.

##### Reward experiment

60 participants completed the task with the following parameters: weak=0.2, W-S ratio=2.5, *p*_1_=0.08. Testing consisted of 600 trials, broken up into two sequential blocks of 300 trials each. For the high-reward block, the bonus increment was ±$0.024/trial. For the no-reward block, participants were not given the opportunity to earn a bonus. Viewing time was 1.5 s. Pre-testing trials consisted of a single set of 100 trials with an “intermediate reward” bonus increment of ±$0.012/trial. The order of the high- and no-reward blocks were counterbalanced across participants. Five more participants were collected to meet the minimum information capacity criterion.

##### Preregistration

The design of these experiments, data collection procedures, and most analyses we presented were preregistered (doi:10.17605/OSF.IO/HQWCY). Specifically, the following analyses and their associated results were preregistered: 1) using bootstrapping to determine if a participant was on an IB bound; 2) measuring the proportion of participants on different bounds; 3) comparisons between participants most consistent with fully optimal behavior and those most consistent with heuristic behavior; 4) within-individual comparisons of information capacity, inference predictiveness, and distance from the IB bound in the learning, decision-time, and reward manipulation experiments; 5) model-fitting and comparison; and 6) simulations of behavior following and not following our softmax solution (equation (3)) for comparison with the ground truth IB bound. We registered other analyses that are not reported here, namely tests to confirm that participants limit their information capacity and that there is variation in information capacity between participants for all experiments and conditions (analyses 3.1 and 3.2 in the preregistration).

#### 4.1.2 Bead-prediction task

The bead-prediction task (a variant of the classic urn task [17]) required an inference about a latent state (the next bead drawn) based on a sequence of past observations (bead draws). One jar contained 80% black beads and 20% white beads (likelihoods), while the other jar contained the opposite percentages. On each trial, participants observed a bead drawn from one of the two jars. Between trials, there was a *h* = 0.01 probability (hazard rate) that the hidden source jar would switch from one to the other. Thus, the temporal sequence of bead draws from the current trial, going backwards in time, was highly informative of the identity of the current source jar (and thus the color of the next bead drawn). Given this statistical structure, the marginal probability distribution of source jar identity was uniform across a large number of trials. 70 participants completed the task.

During the instructions, participants were told about the existence of these two hidden jars. They were also told the exact proportions of beads in each jar and the exact switch rate. They confirmed via a quiz that they learned this information. Finally, they were told that the best way to predict the color of the next bead was to first guess which jar the current bead came from, and then use the hazard rate to predict which jar will be used next. We asked participants to predict the color of the next bead drawn and not the current source jar so that they would receive feedback for their choices without unmasking the current jar (and thus eliminating the need to integrate over the history of observations).

Each participant completed the same sequence of 40 practice trials (40 bead draws), which had been sampled using the same generative statistics (i.e., hidden Markov process parameters) as the main experiment. They then completed two sequential blocks of 500 trials, in which the same exact bead draw sequence was used for each block and participant to standardize IB comparisons across individuals. We analyzed the data from block two as part of the base analysis in Results (Figs. 3 and 4), excluding 6 participants that had an information capacity of less than 0.05 bits (the same exclusion criteria we used for the base horse-prediction experiments). Blocks one and two were compared as the bead-prediction learning experiment in the behavioral adjustment experiments.

Participants received a base payment of $8 for completing the task and a bonus payment equal to 0.02$/percentage point of beads predicted correctly over the 1000 total test trials. The bead-prediction experiment was implemented using jsPsych v7.3.0 [48] and hosted on Pavlovia.

##### Preregistration

This experiment was based on the experiments performed as part of a preregistered study (doi:10.17605/OSF.IO/KZXNQ). In the preregistered experiments, we generated a unique trial set (i.e., bead draw sequence) for each participant using the same generative statistics. We later realized that this sampling variability leads to substantial variability in information capacity, inference predictiveness, information efficiency, and even the position of the IB bounds, obscuring our ability to detect variability driven by true individual differences (Supplemental Fig. A11). Thus, we deemed it necessary to perform a new experiment (not preregistered) in which bead sequences were standardized across participants, as described above. In addition, we had participants complete the same bead sequence twice to evaluate whether participants would have consistent information capacity and inference predictiveness across blocks, all else being equal.

#### 4.1.3 Psychometric functions

We computed psychometric functions for individual participants by grouping trials by unique sets of observations (values of *x* using the full, non-simplified encoding scheme), then determining the probability of making a reference choice for each *x*. We also computed psychometric functions based on non-optimal strategies (see below), such that the x-axis values for each strategy were set to the posterior probability computed using that strategy. We then used a goodness-of-fit measure (KL divergence) to characterize how well data plotted in this way matched a given bound-predicted psychometric function (at the same information capacity as the data).

### 4.2 The Information Bottleneck (IB)

The following is laid out in full detail in [13]. Briefly, let *X* be a discrete signal that contains information about another discrete signal of interest *Y* (*I*(*X*; *Y*) is nonzero), with probability measures *p*(*x*) and *p*(*y*) respectively. Now let *R* be a potentially compressed or lossy representation of the information in *X*. We seek the possibly stochastic mapping between *x* ∈ *X* and *r* ∈ *R*, given by the conditional probability distribution *p*(*r*|*x*), that maximizes the amount of information *R* contains about *Y* (i.e., maximizes *I*(*R*; *Y*), the mutual information between *R* and *Y*). However, suppose that it is costly to increase the amount of information *R* retains from *X* (or in other words *I*(*X*; *R*)). Put another way, we would like to find a balance between lowering this cost of retaining information from *X* and maximizing how much information *R* contains about *Y*. We can formalize this trade-off using the following optimization problem:

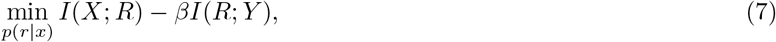

where *I*(*X*; *R*) is the mutual information between *X* and *R*, and *I*(*R*; *Y*) is the mutual information between *R* and *Y*. In general, the conditional probability distribution *p*(*r*|*x*) that solves equation (7) for a given value of *β* satisfies

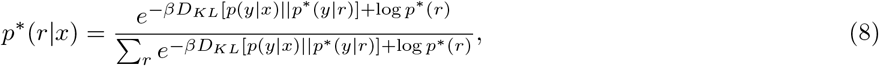

where *p*^∗^ denotes all probability distributions that are derived from *p*^∗^(*r*|*x*) (the ∗ indicates that *p*^∗^(*r*|*x*) satisfies equation (7)), namely

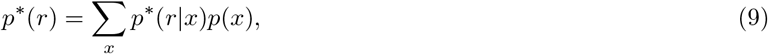

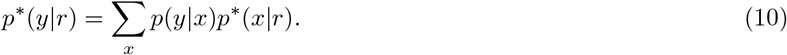

Note that: 1) we must know *p*(*x, y*); and 2) these equations follow from the Markov chain condition *Y* → *X* → *R*, which follows from the setup described. Equation (7) can be solved for a given value of *β* sequentially iterating through equations (8), (9), and (10) until they provably converge.

#### 4.2.1 Application of the IB framework to human inference

Now suppose that *x* ∈ *X* are observations probabilistically related to a latent state *y* ∈ *Y*. A human observer sees the observations *x* and is tasked with inferring the latent state *Y* based on those observations. We treat those inferences as *r* ∈ *R*, which has a natural correspondence to *Y* (e.g., *r* = “observer predicts latent state 1” corresponds to *y* = “actual latent state is 1”). This *R* serves as a proxy for how the observer encodes information from *X* that is relevant for inferring *Y*. An observer who maximizes the amount of information they extract about *Y* from *X* (*I*(*R*; *Y*): inference predictiveness) given the amount of information they encode and use from *X* (*I*(*X*; *R*): information capacity) has choice probabilities *p*(*r*|*x*) that satisfy equation (7) for a unique value of *β*. As in the section above, *p*(*r*|*x*) that satisfy equation (7) for some value of *β* (denoted *p*^∗^(*r*|*x*)) follows equation (8).

For the tasks and inference strategies (fully-optimal or heuristic) we considered, we found that the choice policy solving equation (7) (i.e., satisfying equation (8)) was (a) equally accurate in inferring each latent state, and (b) made each type of error (particular mismatches of *r* and *y*) with the same probability. That is, *p*^∗^(*r* = *i*|*y* = *i*) = *p*^∗^(*r* = *j*|*y* = *j*) for all *i, j* ∈ *Y* (accuracy is the same regardless of latent state), and *p*^∗^(*r* = *s*|*y* = *t*) = *p*^∗^(*r* = *u*|*y* = *v*) for all *s, t, u, v* ∈ *Y* such that *s ≠ t* and *u ≠ v* (error rate is the same regardless of error type). In addition to this “accuracy symmetry”, *p*(*y*) is uniform in our experiments by construction. When these conditions (a) and (b) are met (which we validate in Math Appendix), equation (8) simplifies to the softmax form we reported in Results:

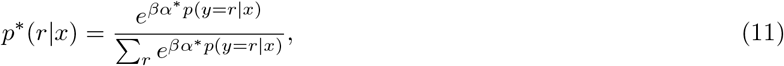

where

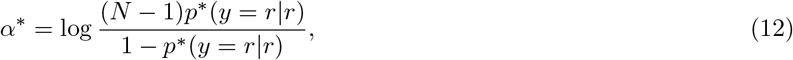

*p*(*y* = *r*|*x*) is the posterior probability of the latent state corresponding to *r*, *p*^∗^(*y* = *r*|*r*) is the probability that the latent state matches the inference that is made, and *N* is the number of unique latent states (see Math Appendix for a derivation). Here *p*^∗^(*y*|*r*) has the same accuracy symmetry as *p*^∗^(*r*|*y*) (in fact, *p*^∗^(*y*|*r*) = *p*^∗^(*r*|*y*)), thus *p*^∗^(*y* = *r*|*r*) refers to the single common accuracy value implied by accuracy symmetry. *p*^∗^(*y* = *r*|*r*) and thus *α*^∗^ are task specific (i.e., they depend on *p*(*x, y*)) and monotonically increase as *β* increases. Consequently, *β*^∗^ = *βα*^∗^ increases monotonically with *β*, establishing a monotonic relationship between information capacity and logistic choice noise (see Math Appendix).

#### 4.2.2 Computing an IB bound for a task

The IB bound tells us the maximum achievable *I*(*R*; *Y*) for each given *I*(*X*; *R*) bits of information from the observations to form the inferences. The default way to compute the IB bound for a task is to define *p*(*x, y*) for that task; select initial values of *p*^∗^(*r*) and *p*^∗^(*y*|*r*); iterate through equations (8), (9), and (10) until *p*^∗^(*r*|*x*) converges; then compute *I*(*X*; *R*) and *I*(*R*; *Y*). This process is repeated over a large range of values of *β*, after which the solution values of *I*(*X*; *R*) and *I*(*R*; *Y*) trace an IB curve (Fig. 2B). When accuracy symmetry holds and *p*(*y*) is uniform, one can analytically compute the bound by plugging in a large range of many values of *β*^∗^ = *βα*^∗^ into equation (11).

However, the finite set of trials used in our experiments can cause slight deviations from the true *p*(*x, y*). Thus, we computed the IB bounds for a given task using the actual trial set in each experiment. Specifically, we used the iterative algorithm described above and the true *p*(*x, y*) to calculate *p*^∗^(*r*|*x*) for each *x* ∈ *X*, and then used these choice probabilities and the empirical *p*(*x, y*) (computed from the finite trial set) to calculate *I*(*X*; *R*) and *I*(*R*; *Y*) over a range of *β* values. Informally, this approach determines the bound for an IB-optimal agent that knows the true generative statistics and is completing a particular finite trial set.

#### 4.2.3 IB analysis of behavioral data

For each task, we first computed the “full” IB bound, which corresponded to the full, non-simplified encoding of the observations *X*. Then, because we had full knowledge of *X*, *R*, and *Y* for each participant, we calculated *I*(*X*; *R*) and *I*(*R*; *Y*) for each. These measures were computed using the empirical distributions of *p*(*x, r*) and *p*(*r, y*) constructed directly from the data (e.g., *p*(*x* = *a, r* = *j*) = *N_x_*_=*a,r*=*j*_*/N_trials_*). We corrected for biases that can arise from estimating these measures from finite trial sets (Supplemental Fig. A12) by using the NSB estimator [49, 50]. We used similar procedures to compute heuristic bounds. As detailed below, these bounds corresponded to various simplified encodings of the observations.

To account for variability in the estimates of *I*(*X*; *R*) and *I*(*R*; *Y*) as a result of choice sampling (Supplemental Fig. A11), we computed 95% confidence intervals on *I*(*R*; *Y*) for each participant using bootstrapping (10000 iterations, trials sampled with replacement the same number of times as number of trials). We considered a participant to be “on” a bound if their confidence intervals overlapped that bound. If the confidence intervals overlapped more than one bound, we considered the participant to be on the closer bound.

We defined “information efficiency”as the inference predictiveness of a participant divided by the predictiveness of the full (or heuristic) IB bound at that participant’s capacity. For our tasks, the information efficiency of the heuristics of interest (one-back for the bead-prediction task, equal-weights for the horse-prediction task) was roughly equal across all information capacities. Thus, it was possible to compare participant information-efficiency values to the average information-efficiency of a heuristic, regardless of capacity.

#### 4.2.4 Within-participant comparisons of IB measures

For the learning, speed-accuracy, and reward experiments, we performed paired-sample Wilcoxon signed-rank tests to determine whether information capacity, inference predictiveness, and/or information efficiency changed across conditions. For the learning experiments, we compared run 1 with run 2. For the speed-accuracy experiment, we compared the “slow” decision-time trials to the “fast” decision-time trials. For the reward experiment, we compared the no reward block to the high reward block. Although we randomized the order of the no and high reward blocks in the reward experiment, we also used a linear mixed effects model to test the effect of reward on the IB measures to further isolate the reward effect from potential block-order or learning effects. In these models, the IB measure (information capacity/inference predictiveness/information efficiency) was the dependent variable while reward condition (no or high) and block number (first or second) were the independent (predictor) variables. Random intercepts were fit for each participant, but an interaction between reward and block number was not included.

We restricted the comparisons of information efficiency to participants who had an information capacity ≥ 0.05 bits across both conditions. This was because the computation of information efficiency involved dividing by a near-zero number when capacity was *<* 0.05 bits, leading to extreme values. Meanwhile, the comparisons of information capacity and inference predictiveness were only restricted to participants who had an information capacity greater than or equal to 0.05 bits in at least one of the two conditions. This was already an exclusion criterion in all weather-prediction experiments, so this did not impact the number of participants in those comparisons. However, this did exclude 2 out of the 70 participants in the bead prediction experiment.

### 4.3 Inference models

#### 4.3.1 Inference models for the horse-prediction task

As captured by equation (3), IB bounds for our tasks correspond to performing exact Bayesian inference of the latent states *Y* using the (potentially simplified) encoding of the observations *X* and then adding varying amounts of logistic choice noise. Thus, we can characterize the noisy optimal inference strategy and the heuristic inference strategies by defining their encoding scheme for *X* and the expression used to calculate *p*(*y*|*x*) for that scheme.

For the horse-prediction task, the noisy optimal inference strategy (corresponding to the full IB bound) uses a full encoding of *X*, which specifies the number of each unique shape type that appears on a given trial. The posterior probabilities *p*(*y*|*x*) are thus computed by summing the true LLRs of the five shapes that appeared (the prior is uniform, so this sum gives the log-posterior odds) and transforming the resulting log-posterior odds (*LPO*, with base 10) into the posterior probability:

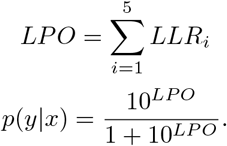

Choice probabilities were then calculated using (3).

The “equal-weighting” heuristic used a simplified encoding in which only the total number of shapes in favor of each latent state were specified without distinguishing between strong and weak shapes. We calculated the posterior probabilities for this strategy using LLRs based on the probabilities that a shape in favor of the green or purple horse would appear.

The “ignore-weak” heuristic used a simplified encoding in which only the strong shapes were encoded if at least one strong shape was present (otherwise the weak shapes were encoded). We calculated the posterior and choice probabilities for this strategy using the same procedure as for the fully optimal inference strategy, but zeroing out the LLRs of any weak shape if a strong shape was present in the trial.

For our model-fitting analyses using for noisy optimal, equal-weighting heuristic, or ignore-weak heuristic strategy, we fixed the LLR to the values computed as described above and fit only *β*^∗^ as a free parameter.

We also considered an inference strategy in which participants weighed the strong and weak shapes differently but deviated from the true relative weighting (the “subjective ratio model”). This strategy used the same algorithm as the noisy optimal strategy, but the relative weighting of the strong and weak shapes varied as a free parameter. For the purposes of model fitting, we fixed the LLR value of the weak shapes to their ground truth value (i.e., the “weak” value used to parameterize the LLRs in these experiments) but allowed the W-S ratio value to vary as a free parameter. *β*^∗^ was also a free parameter. This strategy was identical to the optimal and equal-weighting strategies when the W-S ratio parameter equaled the ground truth value and 1, respectively, and approximated the ignore-weak strategy when the W-S ratio parameter was much higher than the ground truth value.

#### 4.3.2 Inference models for the bead-prediction task

As a discrete hidden Markov process, inference of the latent state (jar) *y_t_* given the sequence of current and past observations (bead draws) *x*_1:*t*_ follows a well-defined form:

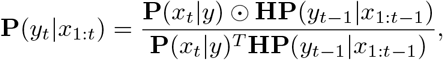

where **P**(*y_t_*|*x*_1:*t*_) is a *N_Y_* × 1 (number of unique latent states *y*) vector giving the posterior probability of each latent state *y* at time point *t* given all observations current and past *x*_1:*t*_, **P**(*x_t_*|*y*) is a *N_Y_* ×1 vector giving the likelihood of the current observation *x_t_* for each *y*, **H** is the *N_Y_* × *N_Y_* transition matrix specifying the probability of transitioning from the latent state at time *t* − 1 (columns) to the latent state at time *t* (rows) (in other words, **H***_i,j_* = *p*(*y_t_* = *i*|*y_t_*_−1_ = *j*)), and **P**(*y_t_*_−1_|*x*_1:*t*−1_) is the vector of posterior probabilities from the previous time point *t* − 1. The true values of these terms reflect the task parameters described above: **P**(*x_t_*|*y*) takes on the likelihood values (0.8 and 0.2) depending on bead and jar identity, and *H* has values of 0.99 on the diagonal (1 − *h*) and 0.01 (*h*) elsewhere. The initial condition, **P**(*y*_0_), is set to the marginal (stationary) distribution of *y*, in this case [0.5,0.5]. The ⊙ operator denotes element-wise matrix multiplication.

This inference process can be understood as updating the posterior probabilities at every time point using the likelihood of the current observation. Specifically, we form a prior for the current time point by filtering the posterior from the previous time step through the transition probabilities (**HP**(*y_t_*_−1_|*x*_1:*t*−1_)), then combine that with the likelihood of the current observation (**P**(*x_t_*|*y*)) to get the posterior for the current time point.

The best possible inference strategy is to use every bead draw observed up to the current trial (*x*_1:*t*_). However, because the ever-increasing sequences of observations on every trial are inherently unique (i.e., each sequence has a different length, namely *t*), it is effectively impossible to construct an empirical distribution *p*(*x, r*). Thus, to define the optimal inference strategy, we restricted *X* to consist of the 7 most recent bead draws observed, which in our data allowed for a reasonably precise estimate of the empirical distribution *p*(*x, r*) given the limited number of trials (500). This strategy computes **P**(*y_t_*|*x_t_*_−6:*t*_), with **P**(*y_t_*_−7_) set to **P**(*y*_0_). Due to the diminishing benefit of integrating over increasing numbers of bead draws (Supplemental Fig. A13), performing Bayesian optimal inference with 7 beads makes the same prediction as the unrestricted Bayesian strategy 98.6% of the time in our task. Thus, this acts as a good approximation of the best-possible inference strategy. For our model-fitting analyses, we fixed **P**(*x_t_*|*y*), **H**, and **P**(*y*_0_) to the ground truth values and fit only *β*^∗^.

We considered a “one-back” heuristic that uses just the most recent bead draw. In general, as the window size decreases, the observation encoding in *X* becomes simpler (fewer beads to encode) while the information efficiency decreases relative to the optimal strategy (Supplemental Fig. A13). This one-back strategy computes **P**(*y_t_*|*x_t_*), which simplifies to **P**(*y_t_*|*x_t_*) = **P**(*x_t_*|*y*) because **P**(*y*_0_) is uniform and **H** is symmetric. Similarly to the above, for our model-fitting analyses we fit only *β*^∗^ for this strategy.

We also considered a strategy (“subjective-hazard model”) in which decision-makers behaved as though they properly integrated over the sequence of bead draws (most recent 7), but had misbeliefs about the statistics of the hidden Markov process. Specifically, we allowed the hazard rate *h* to vary as a free parameter (which determined **H**). We fixed the bead-draw likelihood values to their ground-truth values. Thus, the posterior probabilities were computed in the same manner as for the optimal strategy, just with an **H** that potentially deviated from the ground truth. For model fitting, *h* and *β*^∗^ were fit as free parameters.

#### 4.3.3 Model fitting and comparison

Each inference strategy described above was fit to individual participant choice behavior by maximizing the likelihood of the choices given the model *M* and its parameters *θ*. Specifically, we minimized the negative log-likelihood

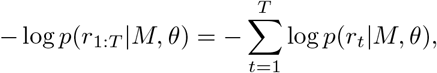

where *r*_1:*T*_ were an individual’s choices and *t* indexed individual trials from the set of *T* total trials. The parameters minimizing this quantity were found using constrained optimization as implemented in the “minimize” function from SciPy with the default algorithm (“L-BFGS-B”). The parameters from the various models were constrained as follows (lower bound, upper bound) - *β*^∗^: (0,None); W-S ratio: (0,None); *h*: (0,1). The default “minimize” options were used for all models except for the subjective-hazard model in the bead prediction task (”eps” set to 1e-19). We additionally performed a grid search over the starting values of the parameters for this model, running “minimize” each time and taking the result with the lowest objective function value (negative sum of the log-likelihoods).

The Bayesian Information Criterion (BIC) [51] was computed for each model fit to evaluate the goodness of fit while accounting for the degrees of freedom each model provided. For the sake of our reported model-comparison results, we took the model with the lowest BIC for a given participant to be the best model of that participant.

## Supplementary information

See Appendix A for supplemental figures. See Appendix B for the math appendix, which includes the fine-grained numerical analyses confirming that equation (3) holds for our inference tasks.

## Acknowledgements

J.A.P. thanks the members of the Gold and Kable labs, along with colleagues in the University of Pennsylvania Computational Neuroscience Initiative, for helpful discussion and feedback on this work. V.B. acknowledges support from the Asha Gupta Chair Professorship at the Centre for Artificial Learning and Intelligence for Biological Research and Education, ICTS-TIFR.

## Declarations

## Data availability

All data presented in this study are available on GitHub at https://github.com/jacobaparker/human-inference-IB.

## Code availability

All code used to process and analyze the data are available on GitHub at https://github.com/jacobaparker/human-inference-IB.

## Author contribution

J.A.P., A.L.S.F., K.L., J.W.K., and J.I.G. conceived and designed the research. J.A.P. derived the solution to the IB problem. J.A.P. and V.B. wrote the mathematical proof of the solution. J.A.P. and A.L.S.F. developed the methods for applying the IB framework to human behavioral data. J.A.P., A.L.S.F., and K.L. implemented the experiments and collected the data. J.A.P. and K.L. performed the data analyses and simulations. All authors contributed to the interpretation of the theoretical and empirical results. J.A.P. wrote the manuscript. All authors revised the manuscript.

## Appendix A Supplemental Figures

**Fig. A1:**
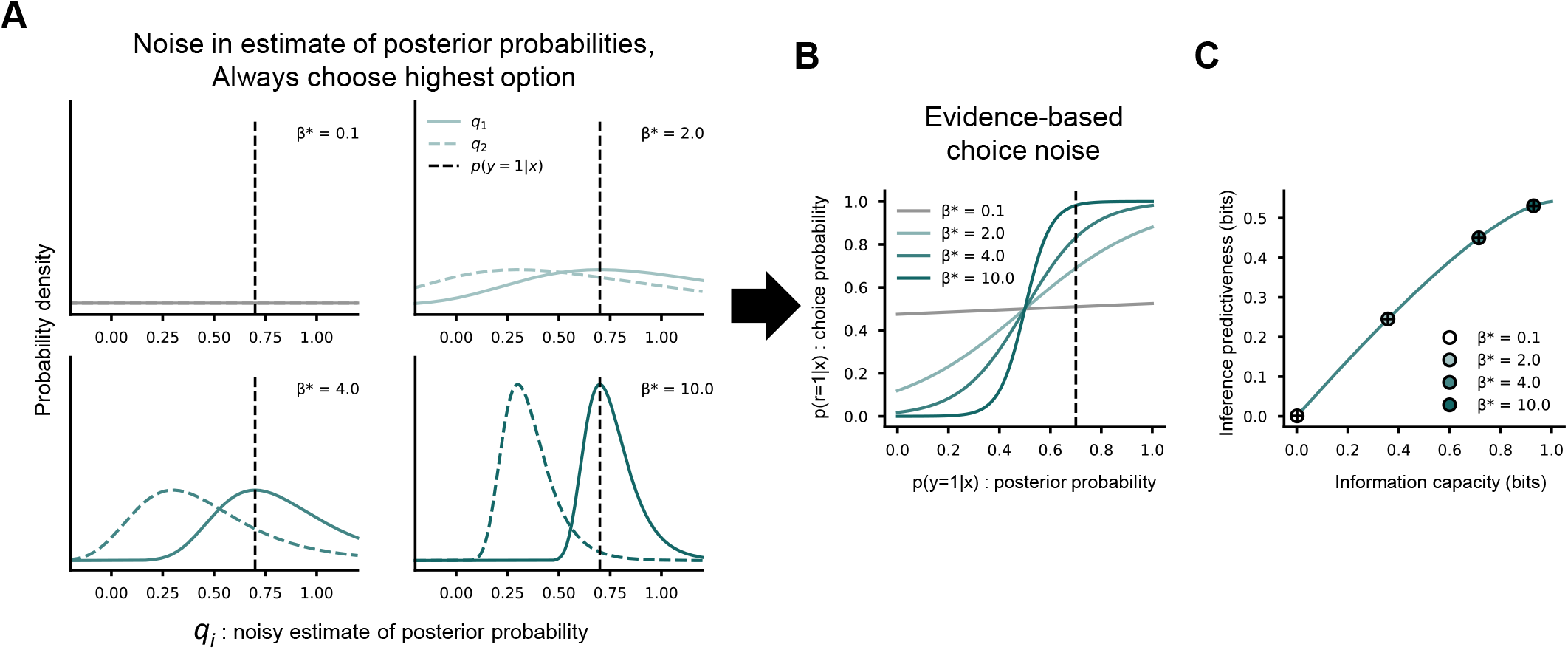
Gumbel noise in estimates of posterior probabilities is equivalent to logistic noise in choices (and is IB-optimal). If we assume that an IB-optimal decision-maker makes a potentially noisy estimate of the posterior probability of each latent state and always chooses the option with the highest estimate (i.e., all choice noise arises from noise in the estimation of the posterior probabilities), these estimates must be drawn from standard Gumbel distributions centered at the true posterior probabilities and scaled by *β*^∗^ (see main text). **(A)** Gumbel distributions corresponding to different values of *β*^∗^ for a particular value of *p*(*y* = 1|*x*) and 1 − *p*(*y* = 1|*x*) (0.7 and 0.3). The probability density function is given by 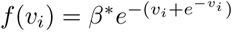, where *v* = *β*^∗^(*q* − *p*(*y* = *i*|*x*)). **(B)** Choosing the highest probability estimate drawn from these Gumbel distributions leads to choice probabilities characterized by logistic choice noise (equation (3)). **(C)** Because this process leads to the same choice probabilities as equation (3), agents using it would be IB-optimal in many of the cases we consider. The points were generated using the softmax solution (equation (3)) while the black crosses were generated using the Gumbel noise process (equations (5) and (6)) for the same values of *β*^∗^.

**Fig. A2:**
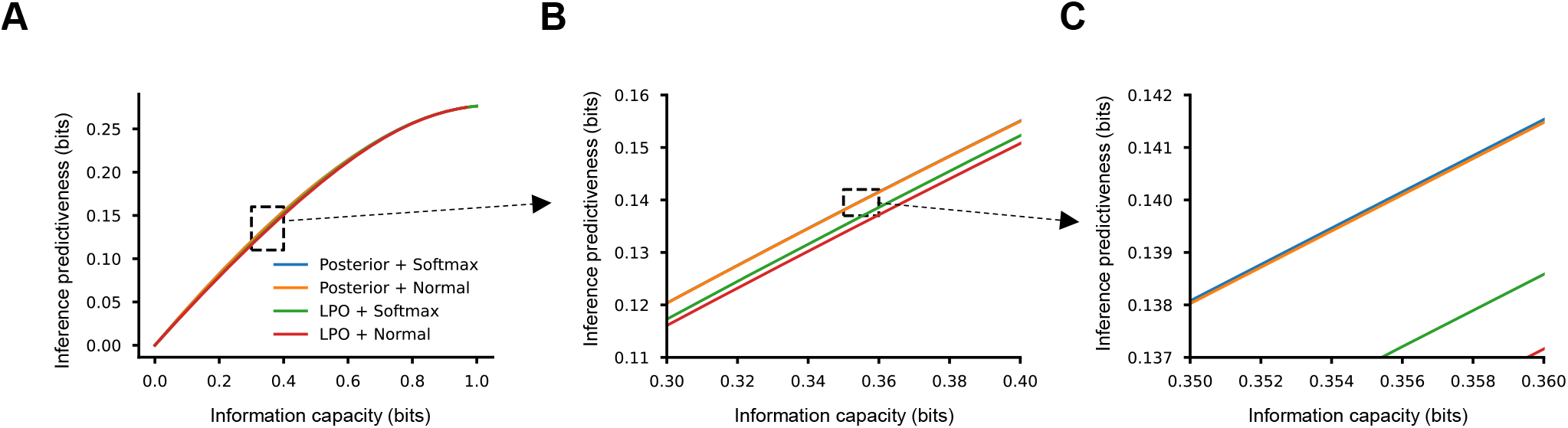
Other common types of evidence-based choice noise are highly information-efficient. **(A–C)** Increasingly zoomed-in plots showing the IB curves computed for common types of evidence-based choice noise. “Posterior” indicates that raw posterior probabilities were used as the evidence, whereas “LPO” indicates that log-posterior odds were used. “Softmax” indicates that the evidence (”posterior” or “LPO”) was transformed into choice probabilities by the standard softmax function, whereas “Normal” indicates that the cumulative normal function was used. The IB curves were computed for each form of choice noise by computing the theoretical choice probabilities over a wide range of noise magnitudes (inverse temperature for logistic, standard deviation for normal) and then *I*(*X*; *R*) and *I*(*R*; *Y*) for each. The statistics of the intermediate W-S ratio horse-prediction experiment (as specified by *p*(*x, y*)) was used for this analysis.

**Fig. A3:**
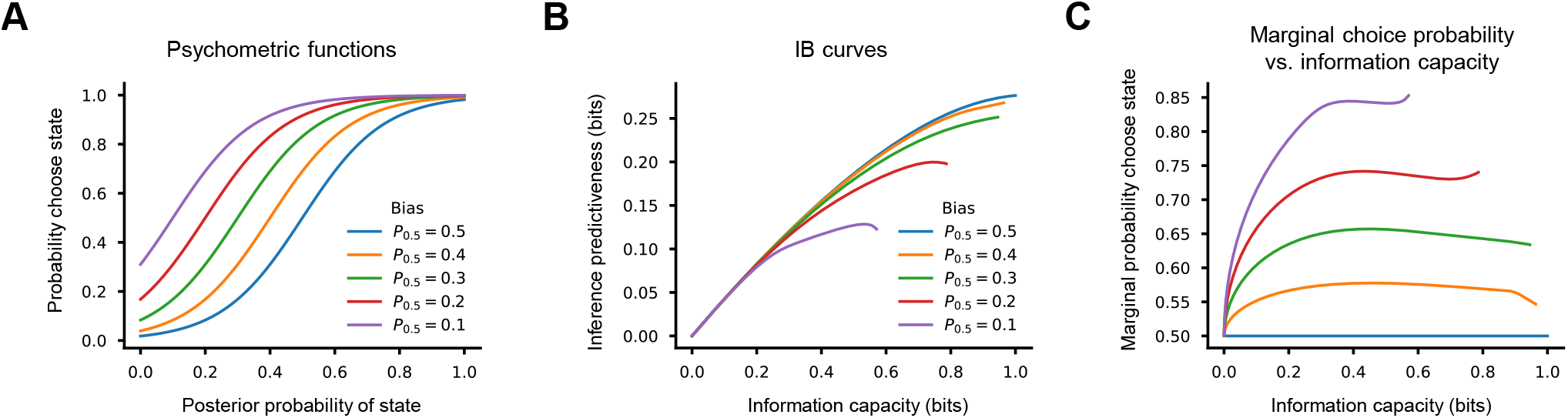
Marginal choice bias (choosing one latent state more often than others, on average) is information-inefficient under the conditions we consider. **(A)** Psychometric functions with increasing amounts of marginal choice bias, generated by translating the standard softmax function. The amount of bias was quantified as *P*_0.5_, the posterior probability at which the choice probability was 0.5 (*P*_0.5_ = 0.5 indicates no bias). **(B)** IB curves corresponding to the amounts of choice bias depicted in A, using the statistics of the intermediate W-S ratio horse-prediction experiment. Because marginal choice bias is information-inefficient at any level of information capacity in this experiment, strategies (points along each curve) with marginal choice bias fall below the IB bound (blue) and are reduced in information capacity for the same degree of logistic noise (value of *β*^∗^). **(C)** Marginal choice probability (probability of choosing a state, on average) as a function of information capacity for each psychometric function. Even when biased choice behavior is described by a translated standard softmax function, the marginal choice probability varies nonlinearly with information capacity.

**Fig. A4:**
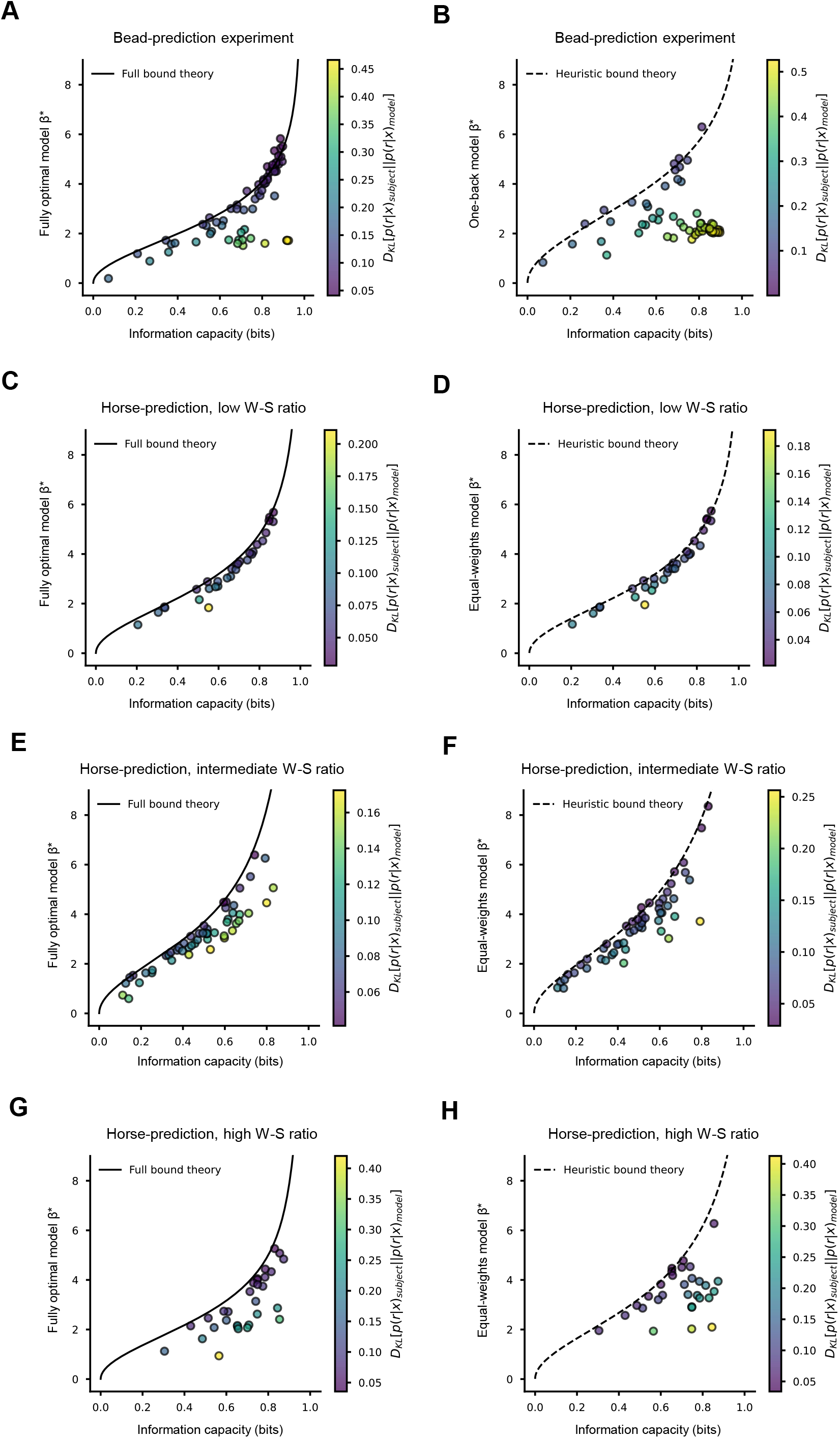
Empirical relationship between choice noise and information capacity converges to theoretical predictions as participant choice behavior more closely matches IB-bound choice behavior. **(A)** Best-fitting value of *β*^∗^ (choice noise magnitude) for the fully optimal model versus information capacity for each participant in block 2 of the bead-prediction experiment. The solid line gives the exact relationship between *β*^∗^ and information capacity for the full IB bound. Participants (points) are colored by the KL divergence between their choice probabilities and the choice probabilities of the best-fitting fully optimal model. As participant behavior more closely matches the fully optimal model (the KL divergence becomes lower), the fitted value of *β*^∗^ (choice noise magnitude) more closely matches the value predicted from the information capacity (using equation 3). **(B)** Same as A, but for the one-back heuristic model. Namely, the plotted values of *β*^∗^, the dotted line, and the KL divergence values are all with respect to the one-back bound and model as opposed to the fully optimal one. **(C), (D)** Same as A and B, but for the low W-S ratio horse-prediction experiment such that C is for the fully optimal model and D is for the equal-weights heuristic model. **(E), (F)** Same as C and D, but for the intermediate W-S ratio horse-prediction experiment. **(G), (H)** Same as C and D, but for the high W-S ratio horse-prediction experiment. In all cases, the observed empirical relationship between logistic choice noise (fitted *β*^∗^) and information capacity approaches the theoretical prediction for a given bound as participant behavior more closely matches the corresponding model.

**Fig. A5:**
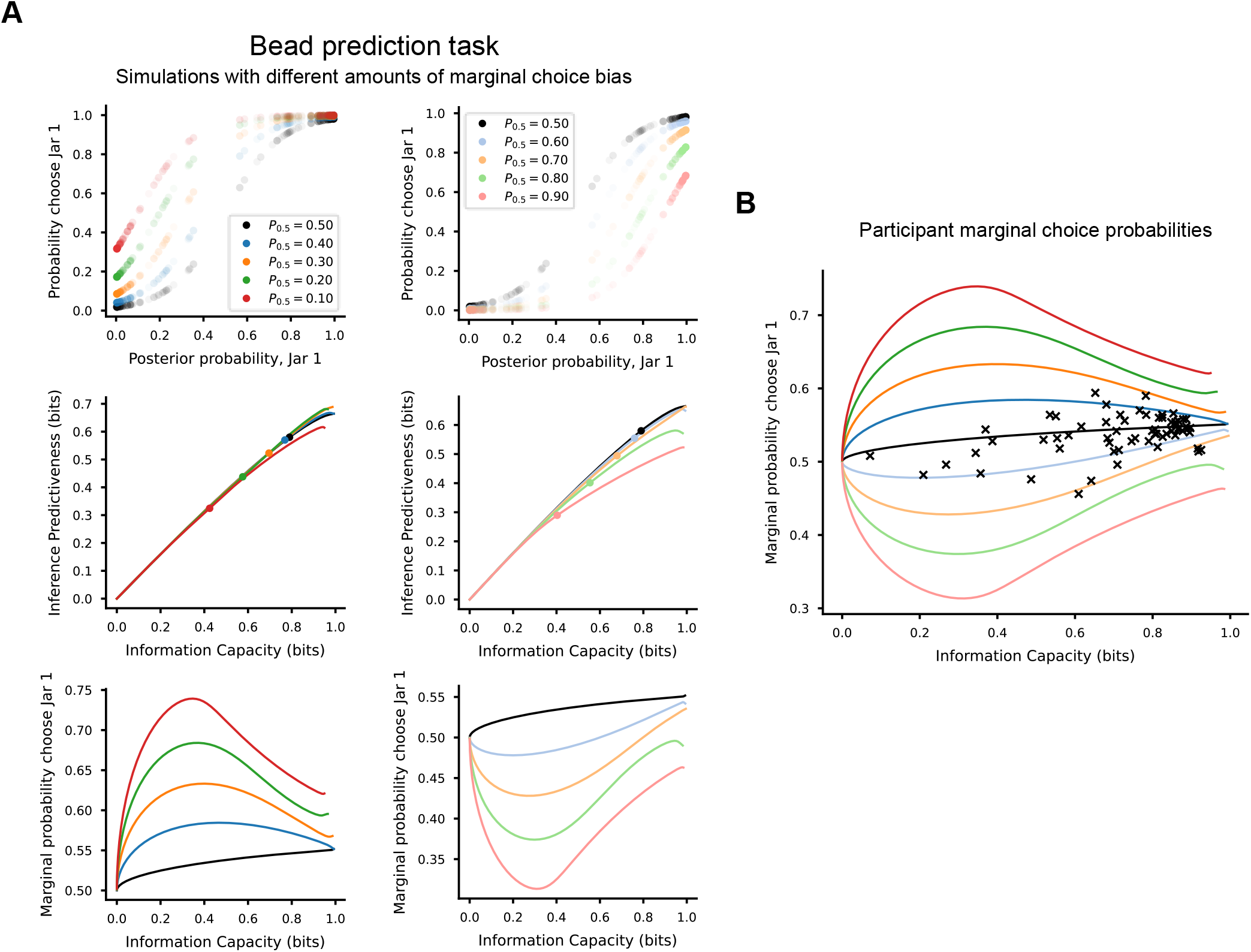
Participant marginal choice probabilities in block 2 of the bead-prediction experiment. **(A)** Simulating the effects of marginal choice bias on performance for the bead-prediction task. (Top row) Psychometric functions for different amounts of marginal choice bias, generated by translating the standard softmax function. Degree of marginal choice bias is given by *P*_0.5_, the posterior probability (fully optimal strategy) at which the choice probability is 0.5. Each point indicates a specific trial type (bead draw sequence); shading indicates the relative frequency in the trial set. (Middle row) IB curves corresponding to the amounts of marginal choice bias depicted above. The black curves (corresponding to no bias) on each plot are the fully optimal IB-bound. Some of the positive bias curves are above the bound (left middle) because for this trial set, Jar 1 occurred slightly more frequently than Jar 2. The points denote the information capacity and inference predictiveness corresponding to the psychometric functions in the panels above. Notably, even with equal psychometric slopes (equal values of *β*^∗^), increasing bias reduces capacity (in addition to falling further from the IB bound). (Bottom row) Marginal choice probability as a function of information capacity for each psychometric function. The non-biased strategy (black curve) increases slightly with capacity because Jar 1 occurs slightly more frequently than Jar 2. Given the complex relationship between degree of true marginal choice bias, observed marginal choice probability, and information capacity (as displayed here), it is important to take into account information capacity when evaluating participant marginal choice probabilities. **(B)** Participant marginal choice probabilities (X’s) plotted as a function of information capacity. As a point of reference, the curves from the bottom row of A are also plotted. All subjects fall in between (or near) the orange and light-orange curves, which is an amount of marginal choice bias that would cause a decision-maker to appear nearly indistinguishable from the full IB bound (middle row, A).

**Fig. A6:**
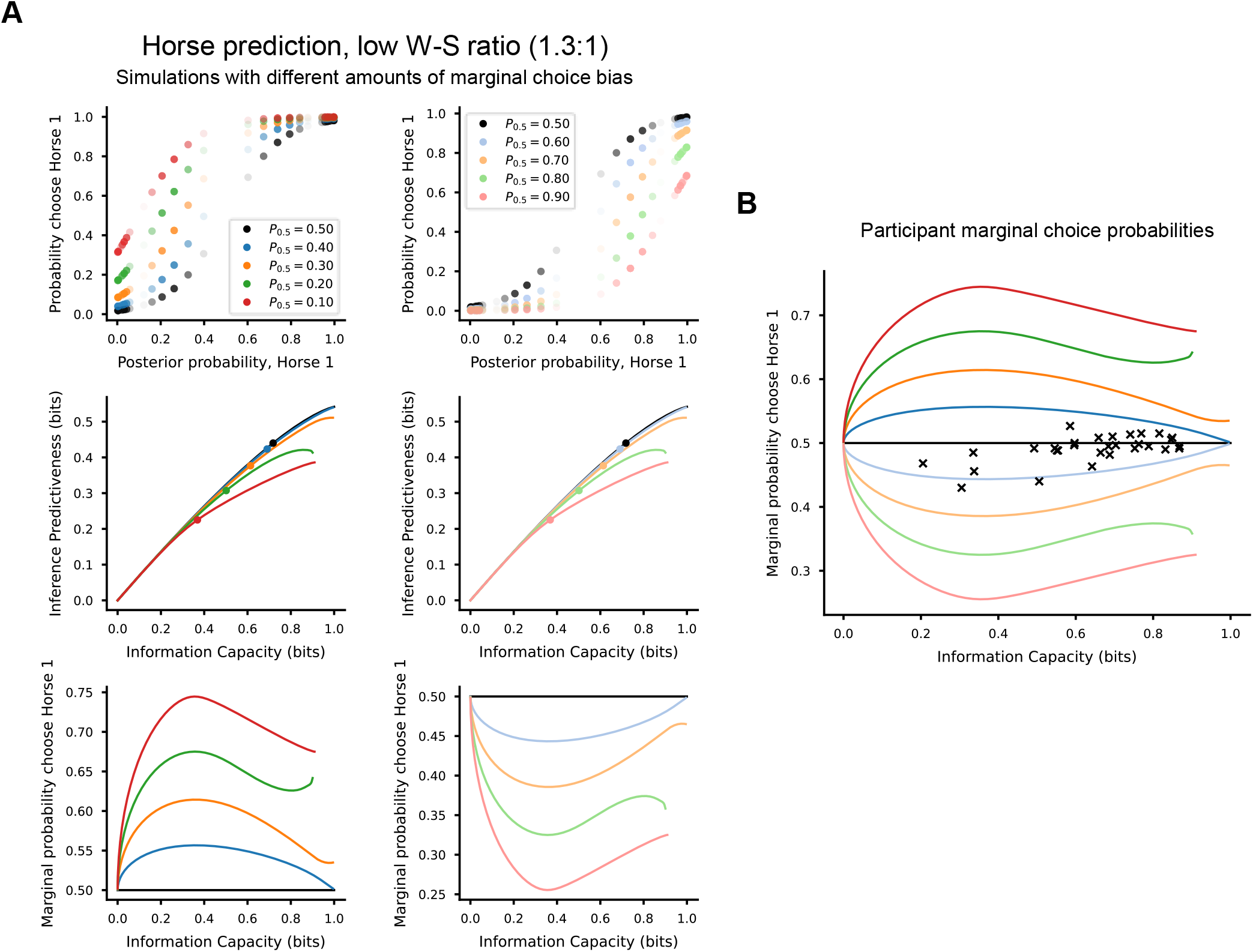
Participant marginal choice probabilities in the low W-S ratio horse prediction experiment. **(A), (B)** Same as Figure A5, but for the low W-S ratio horse-prediction experiment. All subjects fall in-between the orange and light-orange curves, which is an amount of marginal choice bias that would cause a decision-maker to appear nearly indistinguishable from the full IB bound (middle row, A).

**Fig. A7:**
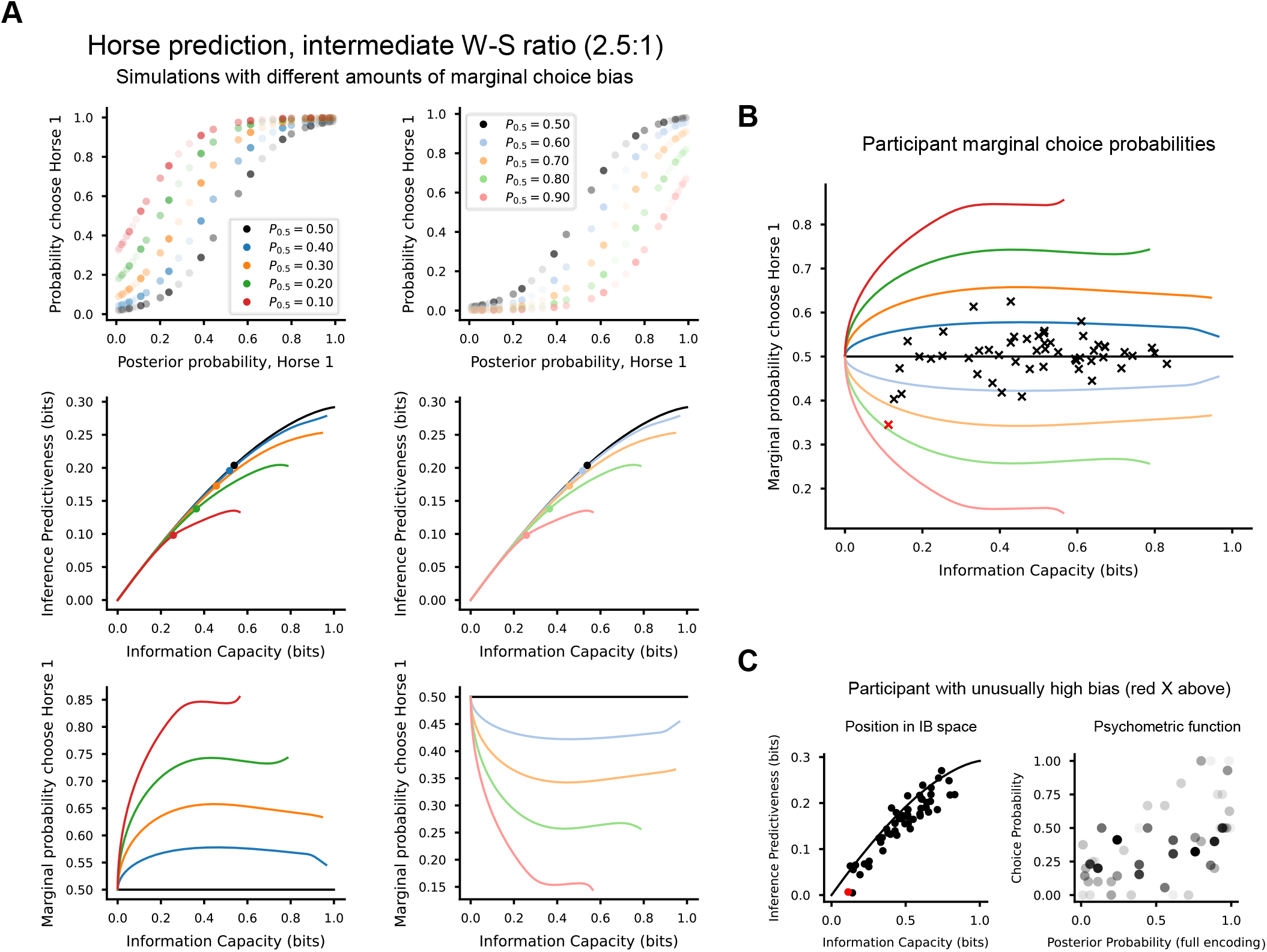
Participant marginal choice probabilities in the intermediate W-S ratio horse prediction experiment. **(A), (B)** Same as Figure A5, but for the intermediate W-S ratio horse-prediction experiment. **(C)** The position in IB space (left) and the psychometric function (right; see Fig. 3, 5 for details on construction) of a participant with an unusually high amount of marginal choice bias (red X in B, red circle in C). All other subjects fall in between the orange and light-orange curves, which is an amount of marginal choice bias that would cause a decision-maker to appear nearly indistinguishable from the full IB bound (middle row, A).

**Fig. A8:**
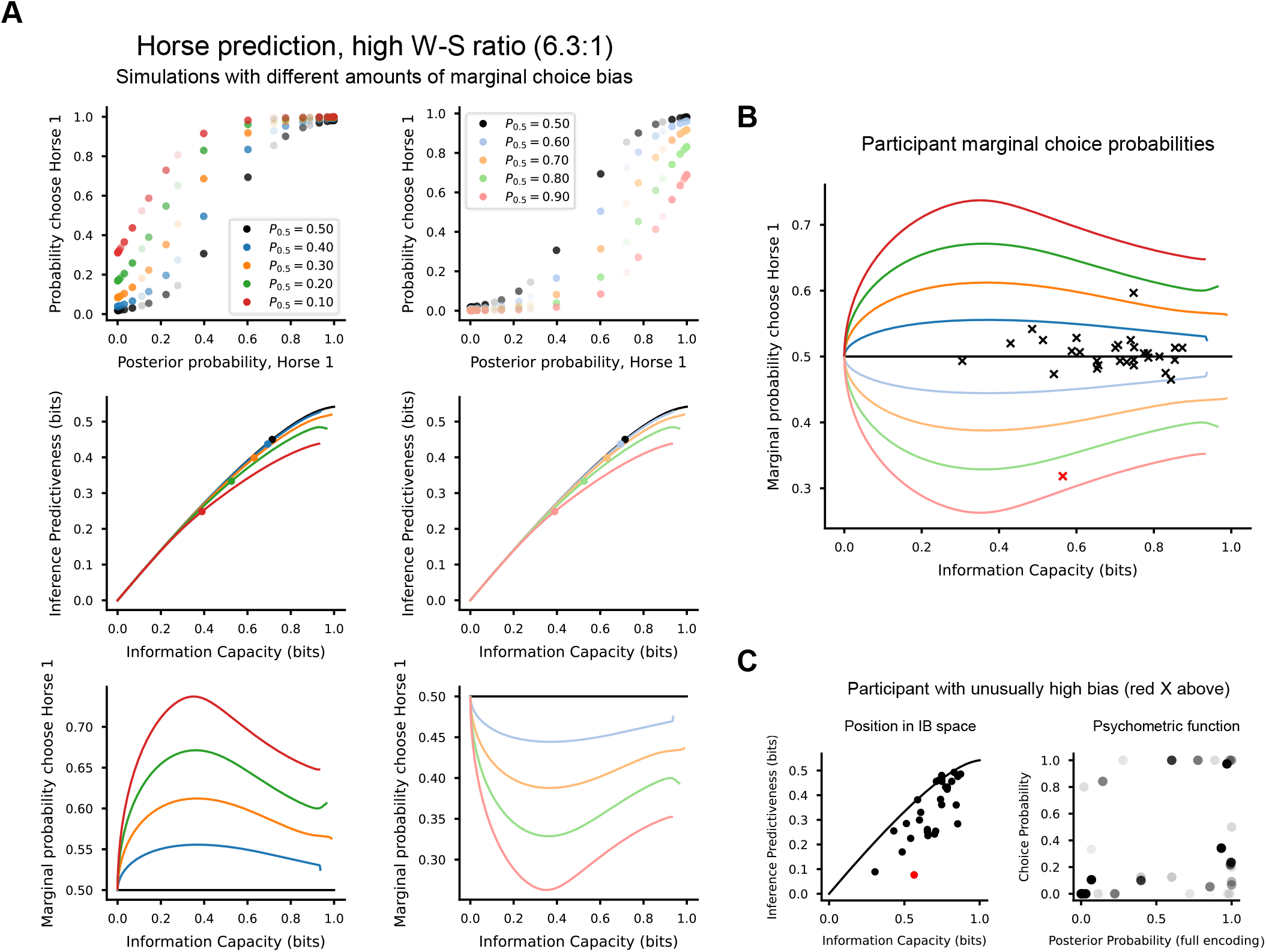
Participant marginal choice probabilities in the high W-S ratio horse prediction experiment. **(A), (B)** Same as Figure A5, but for the high W-S ratio horse-prediction experiment. **(C)** The position in IB space (left) and the psychometric function (right; see Fig. 3, 5 for details on construction) of a participant with an unusually high amount of marginal choice bias (red X in B, red circle in C). All other subjects fall in between (or near) the orange and light-orange curves, which is an amount of marginal choice bias that would cause a decision-maker to appear nearly indistinguishable from the full IB bound (middle row, A).

**Fig. A9:**
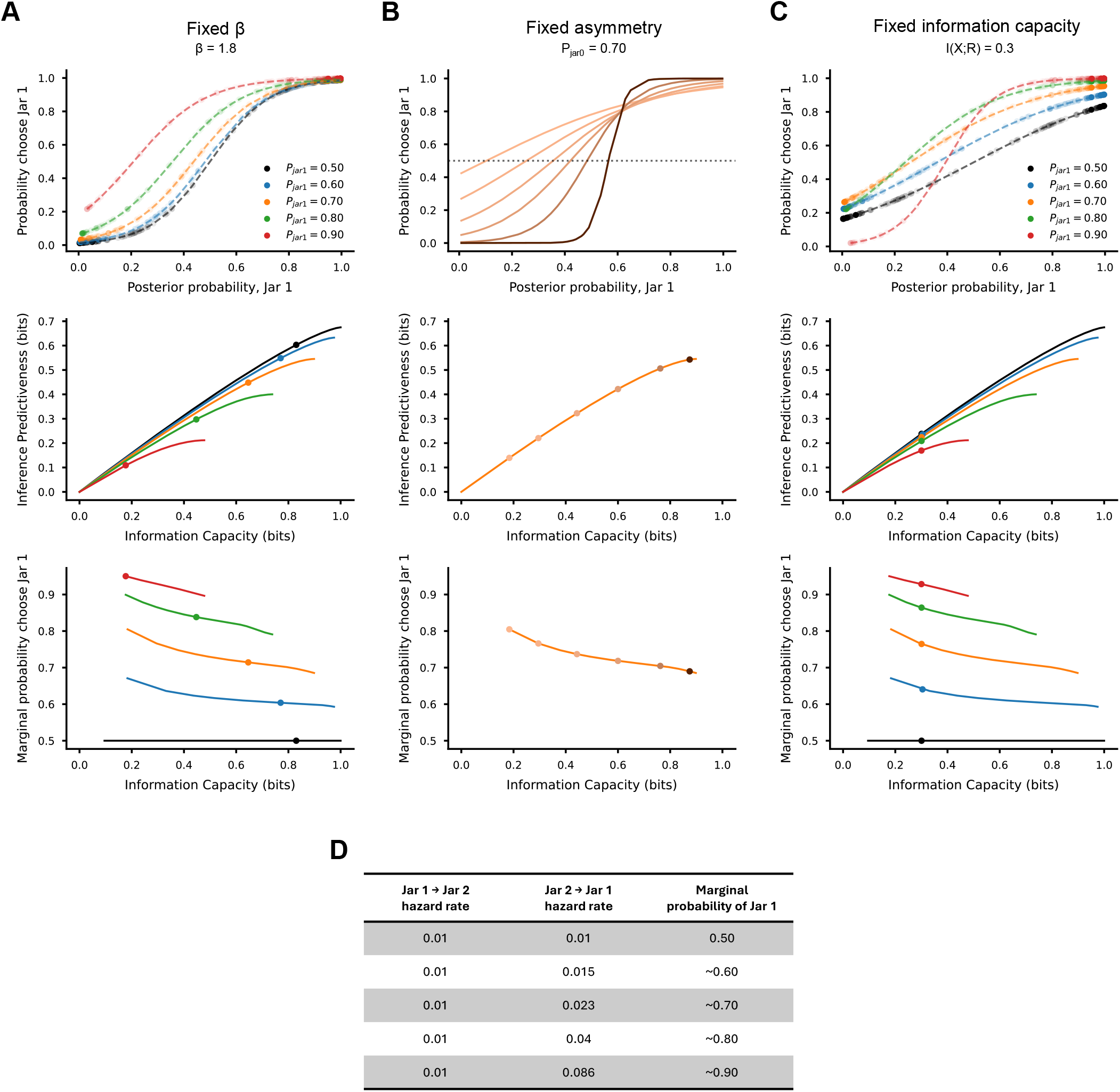
IB bounds and IB-optimal choice behavior for different marginal latent-state distributions (*p*(*y*)) for the bead-prediction experiment. In our experiments, the marginal latent state distribution *p*(*y*) was always uniform. Here, we take a look at the characteristics of IB solutions and bounds for simulated versions of the bead prediction task for varying degrees of asymmetry in *p*(*y*). **(A)** Analyses with a fixed value of *β*. (Top) Psychometric functions describing IB-optimal behavior for different marginal latent state distributions (all for the full IB bound). Individual points denote specific trial types (bead draw sequences); shading indicates relative frequency (which changes as *p*(*y*) changes). The dashed line connects the points, for visualization. (Middle) IB bounds for each marginal latent state distribution. Points indicate the position corresponding to each psychometric function above. (Bottom) Marginal choice probability as a function of information capacity for each marginal latent-state distribution. The point again indicates the position corresponding to each psychometric function. **(B)** Same as A, but varying *β* with a fixed marginal latent state distribution, in this case (*P_jar_*_1_*, P_jar_*_2_) = (0.7, 0.3). **(C)** Same as A, but fixing information capacity and varying the marginal distribution. As expected, IB-optimal behavior (as characterized by the psychometric functions in the top row) departs from the standard softmax once *p*(*y*) becomes non-uniform. Furthermore, an asymmetric *p*(*y*) induces an inverse relationship between marginal choice probability and information capacity such that IB-optimal choice behavior becomes increasingly biased toward the more frequent *y* as capacity decreases. **(D)** The marginal latent-state distribution was manipulated by altering the hazard rate for Jar 2 to Jar 1 transitions. The table gives the exact hazard rates used and the marginal probability of jar 1 they corresponded to. The latter can be calculated by normalizing the first eigenvector of the transition matrix (constructed using the given hazard rates, see Methods), or by raising the transition matrix to large power and then multiplying by a vector of probabilities.

**Fig. A10:**
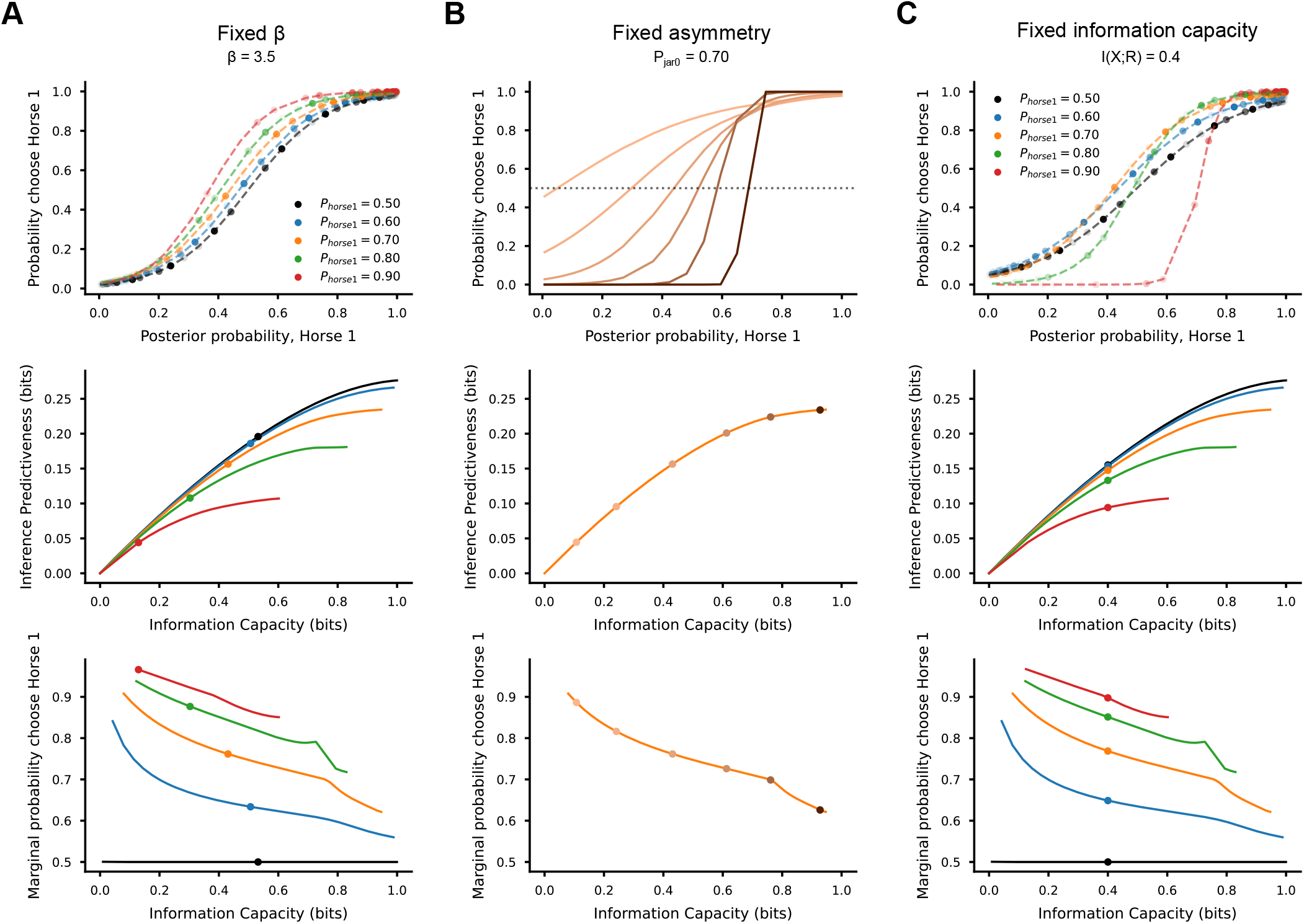
IB bounds and IB-optimal choice behavior for different marginal latent state distributions (*p*(*Y*)) for the horse prediction experiment (intermediate W-S ratio). **(A), (B), (C)** Same as A, B, and C in Figure A9. As expected, IB-optimal behavior (as characterized by the psychometric functions in the top row) departs from the standard softmax once *p*(*y*) becomes non-uniform. Furthermore, an asymmetric *p*(*y*) induces an inverse relationship between marginal choice probability and information capacity such that IB-optimal choice behavior becomes increasingly biased toward the more frequent *y* as capacity decreases.

**Fig. A11:**
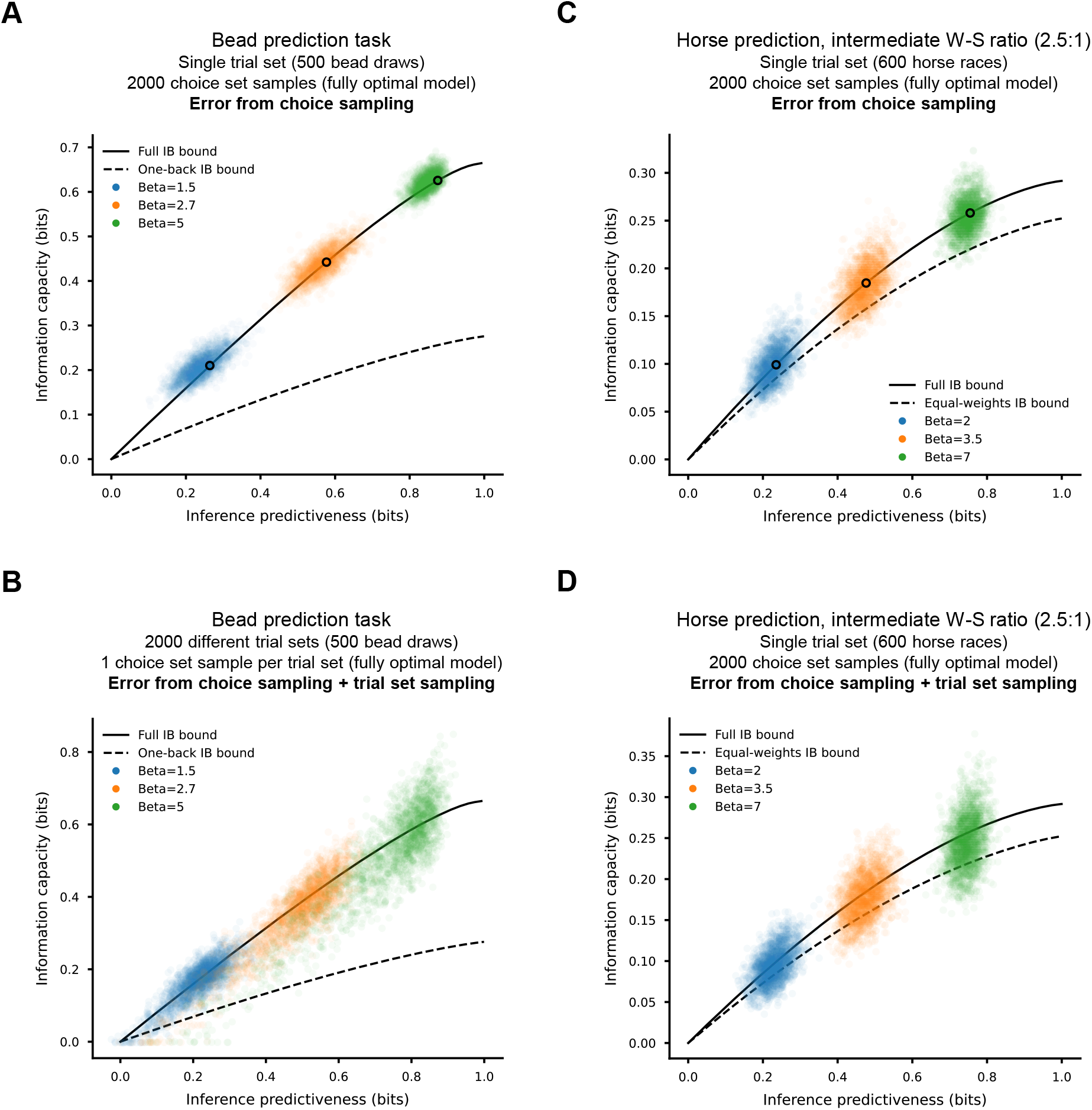
Measurement error in IB measures that result from choice sampling and trial-set sampling. **(A)** Variability in measured information capacity and inference predictiveness that result from choice sampling in the bead-prediction experiment. Each point cloud (corresponding to particular values of *β*^∗^, as indicated) shows the estimated information capacity and inference predictiveness for 2000 choice sets sampled from the fully optimal model on the actual set of 500 trials used in the experiment. The black circle gives the theoretical value of information capacity and inference predictiveness corresponding to the ground truth choice probabilities. **(B)** Same as A, except a different set of 500 trials is used for each of the 2000 sampled choice sets. Thus, measurement variability reflects not only choice sampling variability, but also variability from sampling a set of trials. **(C), (D)** Same as A and B, but for the intermediate W-S ratio horse-prediction experiment.

**Fig. A12:**
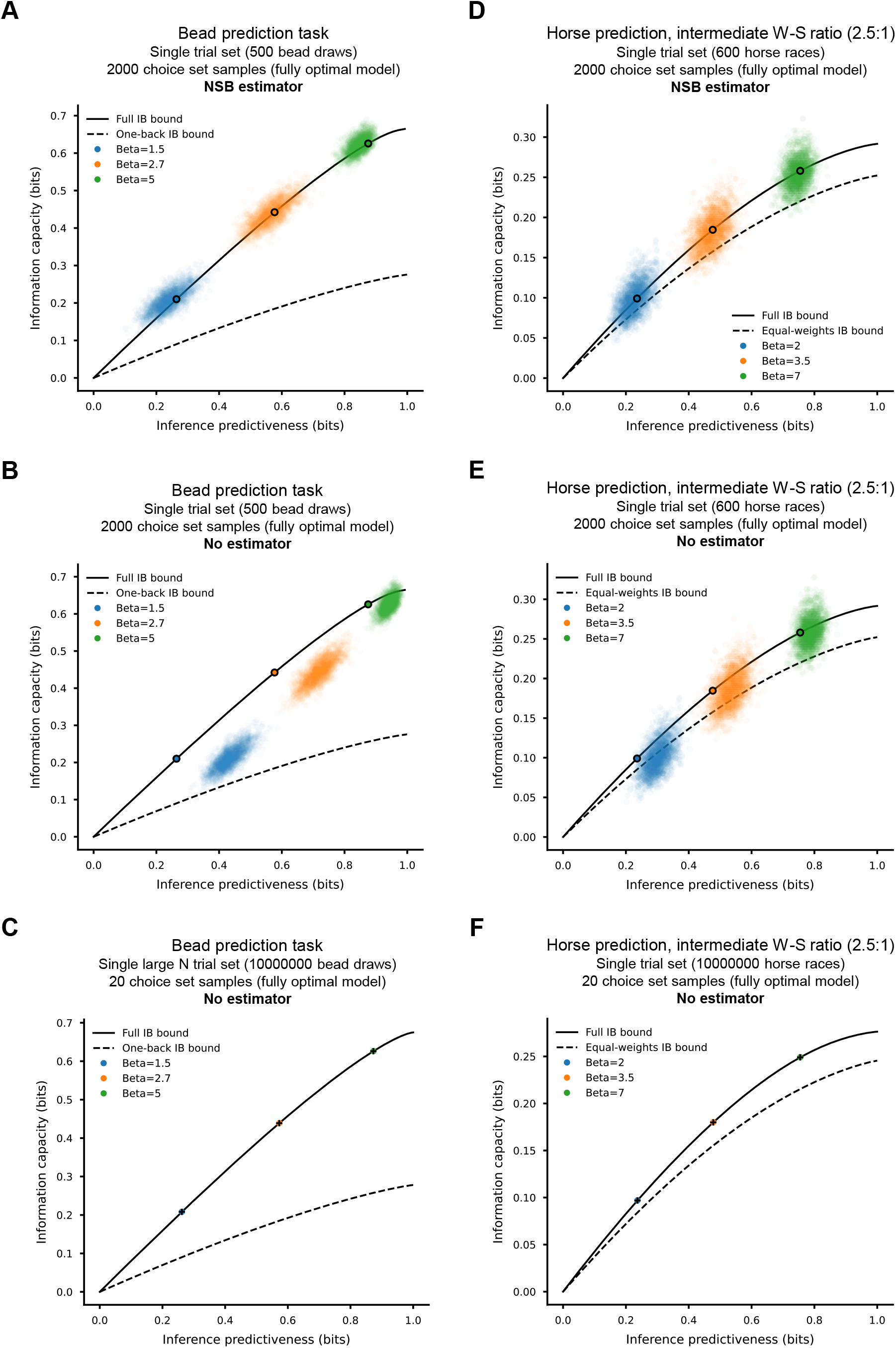
An estimator is required to correct for biases in the estimation of IB measures on human choice data. Because of the finite number of trials obtained in human behavioral experiments, using the standard formula for mutual information results in substantial biases in estimated IB measures. **(A)** Each point cloud (corresponding to different values of *β*^∗^) shows the estimated information capacity and inference predictiveness for 2000 choice sets sampled from the fully optimal model on the actual set of 500 trials used in the experiment. Crucially, the NSB estimator [49] was used to estimate information capacity and inference predictiveness. The black circle gives the theoretical value of information capacity and inference predictiveness corresponding to the ground truth choice probabilities. **(B)** Same as A, but this time using no estimator (the standard formula for mutual information) to estimate information capacity and inference predictiveness. Even though the fully-optimal model was used to generate choice sets, all are substantially shifted in IB space relative to the theoretical value. **(C)** Same as B, but this time for a single very large trial set (10000000 bead draws). In this regime, information capacity and inference predictiveness converge to the theoretical values for the ground truth *p*(*x, y*) (black crosses). This result suggests that the bias observed in B is a consequence of the relatively finite number of trials. **(D), (E), (F)** Same as A, B, and C but for the intermediate W-S ratio horse-prediction experiment.

**Fig. A13:**
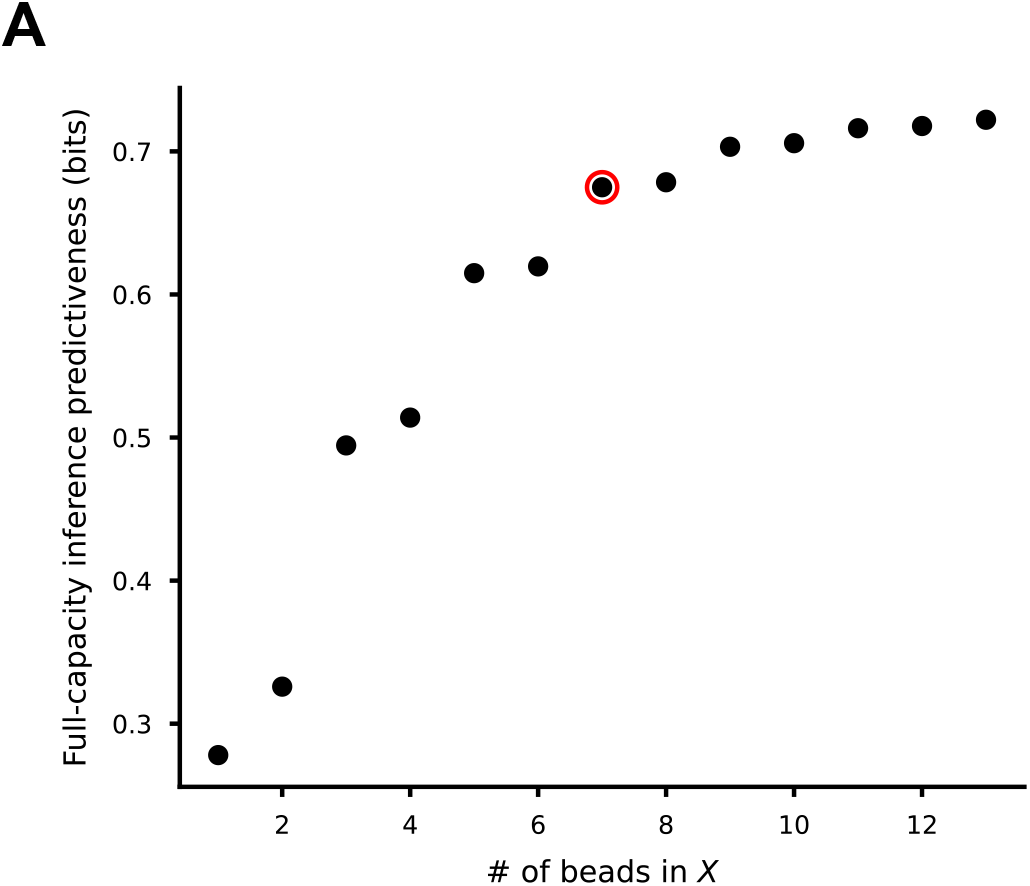
Inference predictiveness at maximum information capacity as a function of number of bead draws included in *X*. Computed using the true generative statistics (*p*(*x, y*)) of the bead-prediction task. Red circle indicates the number of beads we used to approximate the optimal strategy (using all previous bead draws to make inferences). As more bead draws are included in *X*, the marginal benefit of each new bead included quickly vanishes. Thus, the seven most recent bead draws captured most of the predictive information contained within the entire history of bead draws.

## Appendix B Math Appendix

As discussed in Results and Methods, we found that the solution to the IB problem obeys the classic softmax expression (Eq. 3) for the inference tasks, and for all full and heuristic strategies, that we considered. In this appendix, we describe the conditions that must be satisfied for the softmax solution to hold and validate that they are satisfied for the tasks and strategies that we discussed.

### B.1 Derivation of the softmax IB solution

Let *X* be discrete observations probabilistically related to a latent state *Y* such that *I*(*X*; *Y*) *>* 0. Let *R* be the potentially stochastic inferences a decision-maker makes about *Y* after observing *X*. In other words, *r* ∈ *R* is what the decision-maker believes *Y* to be after they have observed some *x* ∈ *X*. Given this definition, it follows that *R* and *Y* have the same number of elements with a natural correspondence (e.g., *r* =“infer jar 1” corresponds to *y* =“jar 1”). If it is costly to encode and use information from the observations to form inferences, a decision-maker may seek to limit (or compress) this information. In formal terms, they seek to limit the information capacity *I*(*X*; *R*). However, assume the decision-maker wants to achieve the best performance possible for a given information capacity. One measure of performance is inference predictiveness *I*(*R*; *Y*), which quantifies the amount of information the inferences contain about the latent state. We can solve the IB optimization problem to find the choice probabilities *p*(*r*|*x*) that allow a decision-maker to maximize inference predictiveness given a limited information capacity:

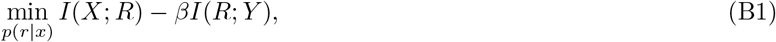

where *β* ∈ [0, ∞) controls the relative importance of performance and information capacity. In general, the values of *p*(*r*|*x*) that satisfy (B1) for a given *β* obey:

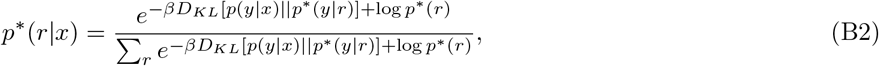

where *D_KL_*is the Kullback-Leibler (KL) divergence, and *p*^∗^ denotes all probability distributions that are derived from *p*^∗^(*r*|*x*), which satisfies (B1), namely

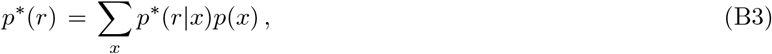

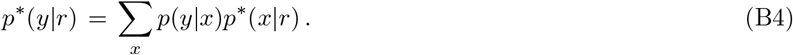

**Theorem 1** *If (1) p(y) is uniform, (2) p^∗^(r = i|y = i) = p^∗^(r = j|y = j) for all i, j ∈ Y, and (3) p^∗^(r = s|y = t) = p^∗^(r = u|y = v) for all s, t, u, v ∈ Y such that s≠ t and u≠ v, then*

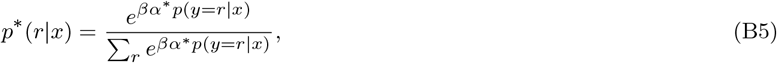

*where*

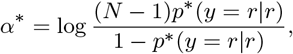

*and p*^∗^(*y* = *r*|*r*) *is shorthand for p*^∗^(*y* = *i*|*r* = *i*)*, which is the same for all values i* ∈ *Y, and N is the number of elements in Y and R (i.e., the number of latent states)*.

The conditions stated in Theorem 1 imply that the IB-optimal choice strategy that solves (B1) has what we will term *accuracy symmetry*: it is equally accurate in inferring each latent state (i.e., the probability of making a correct inference does not depend on latent state identity), and makes all types of errors (particular mismatches of *r* and *y*, e.g., *r* = 1 when *y* = 2) with the same probability.

*Proof* To derive (equation B5), we will first show that *p*^∗^(*r*) is uniform and that *p*^∗^(*y*|*r*) has accuracy symmetry given the conditions of the Theorem. Because *p*(*y*) is assumed uniform, computing *p*^∗^(*r, y*) involves multiplying every *p*^∗^(*r*|*y*) by the same constant value, *p*(*y*) = 1*/N*. This operation does not change the equalities present in *p*^∗^(*r*|*y*), which reflect accuracy symmetry as discussed above. Hence *p*^∗^(*r*) is also uniform because *p*^∗^(*r*) = Σ*_y_ p*^∗^(*r, y*). That is, no matter the value of *r*, the terms of the sum over *y* are the same. Specifically, one term corresponds to the single “accuracy” rate (when *r* = *y*), and the *N* − 1 remaining terms correspond to the single “inaccuracy” rate (any mismatch of *r* and *y*).

Given that *p*^∗^(*r*) is uniform and *p*^∗^(*r, y*) has accuracy symmetry, it follows that *p*^∗^(*y*|*r*) also has accuracy symmetry. Specifically, to compute *p*^∗^(*y*|*r*), we simply divide *p*^∗^(*r, y*) by the same constant value *p*^∗^(*r*) = 1*/N* in every case, thus conserving the accuracy symmetry in *p*^∗^(*r, y*) and in *p*^∗^(*r*|*y*). Thus, *p*^∗^(*y* = *i*|*r* = *i*) = *p*^∗^(*y* = *j*|*r* = *j*) for all *i, j* ∈ *Y* and *p*^∗^(*y* = *t*|*r* = *s*) = *p*^∗^(*y* = *v*|*r* = *u*) for all *s, t, u, v* ∈ *Y* such that *s ≠ t* and *u ≠ v*.

Below we use *p*^∗^(*y* = *r*|*r*) to refer to the unique “accuracy” value of *p*^∗^(*y* = *i*|*r* = *i*). Further, we use *p*^∗^(*y ≠ r*|*r*) to refer to the unique error probability *p*^∗^(*y* = *i*|*r* = *j*) for any *i ≠ j*. Note that the total probability of having a mismatch, 1 − *p*(*y* = *r*|*r*), is the sum of all possible mismatches for a given value of *r*, Σ*_y_*_̸=*r*_ *p*^∗^(*y*|*r*) = (*N* − 1)*p*^∗^(*y ≠ r*|*r*), since there are *N* − 1 mismatches for a single value of *r*.

With the properties of *p*^∗^(*r*) and *p*^∗^(*y*|*r*) established, we can now derive (equation B1). First notice that the KL divergences in the numerator and denominator of (B2) take the form *D_KL_*[*p*(*y*|*x*||*p*^∗^(*y*|*r*)] = Σ_y_*p*(*y*|*x*) log (*p*(*y*|*x*)*/p*^∗^(*y*|*r*)) = Σ*_y_ p*(*y*|*x*) log *p*(*y*|*x*) − Σ*_y_ p*(*y*|*x*) log *p*^∗^(*y*|*r*). The fixed value Σ*_y_ p*(*y*|*x*) log *p*(*y*|*x*) cancels between the numerator and denominator of (B2), giving

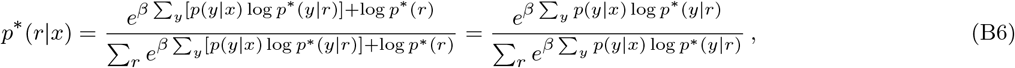

where we used the uniformity of *p*^∗^(*r*) to get the second expression by canceling factors in the numerator and denominator.

The exponents in the numerator and denominator can be simplified as follows. First we split the sum on *y* into *y* = *r* and *N* − 1 terms with *y ≠ r*:

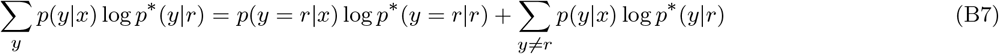

Next observe that by the accuracy symmetry *p*^∗^(*y*|*r*) is independent of *y* and *r* for all *y ≠ r*. So we can pull it out of the sum in the second term to get

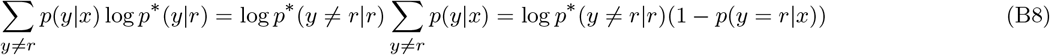

where we used the fact that Σ*_y_ p*(*y*|*x*) = 1. Combining (B7) and (B8) gives

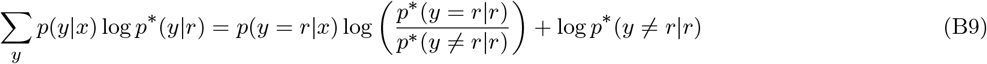

Finally note that

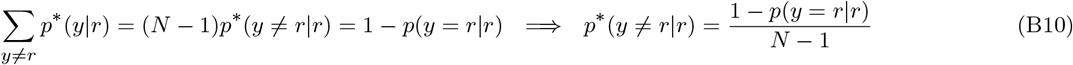

where we used the accuracy symmetry again to get the first equality. Now we apply (B10) in (B9), and insert the result into the exponents in the (B6). After using the accuracy symmetry once more to cancel a common factor of exp(*β* log *p*^∗^(*y ≠ r*|*r*) between the numerator and denominator, we find the result

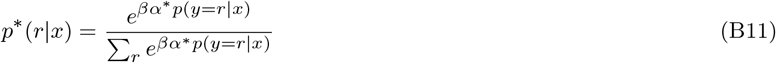

where

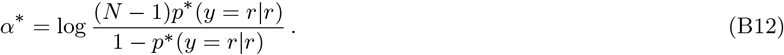

*α*^∗^ is a monotonically increasing function of *β*, which implies that each value of *β* corresponds to a unique value of *βα*^∗^. Thus, increasing the value of *βα*^∗^ (≥ 0) corresponds to moving upward and to the right along the IB bound. Put another way, information capacity is a monotonically increasing function of *βα*^∗^, and thus the degree of evidence-based logistic choice noise corresponds to the information capacity of inference.

We can confirm these relationships by showing that *I*(*R*; *Y*) is a monotonically increasing function of *p*^∗^(*y* = *r*|*r*). We start by expressing *I*(*R*; *Y*) in terms of *p*^∗^(*y* = *r*|*r*):

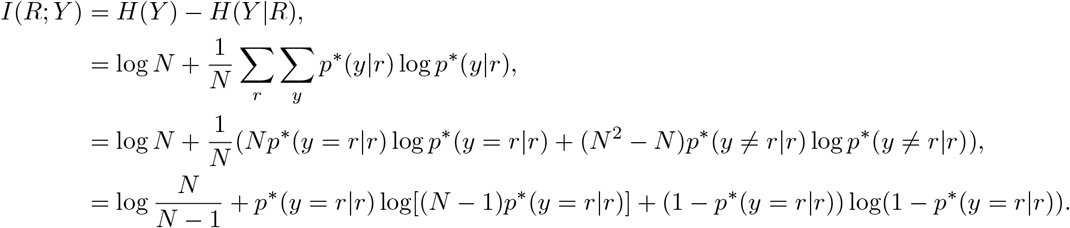

To get the second line we used the definition of entropy and conditional entropy, and the uniform probability of *y* ∈ *Y*. To get the third line we split the sum on *y* into *y* = *r* and *y ≠ r*, and used accuracy symmetry to recognize that for each of the *N* values of *r* and the *N* − 1 values of *y ≠ r* the summand will be the same, i.e., *p*^∗^(*y ≠ r*|*r*) log *p*^∗^(*y ≠ r*|*r*)), leading to a factor of *N* (*N* − 1) from the sums on *r* and *y ≠ r*. To get the last line we used (B10) to write *p*^∗^(*y ≠ r*|*r*) in terms of *p*^∗^(*y* = *r*|*r*) and and then rearranged the resulting expression. Taking the derivative of *I*(*R*; *Y*) with respect to *p*^∗^(*y* = *r*|*r*), we find

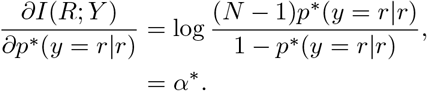

In the range *α*^∗^ ≥ 0, we find that *I*(*R*; *Y*) = 0 when *α*^∗^ = 0 (which corresponds to *p*^∗^(*y* = *r*|*r*) = 1*/N*). When *α*^∗^ *>* 0, it is clear that *∂I*(*R*; *Y*)*/∂p*^∗^(*y* = *r*|*r*) *>* 0, which implies that *I*(*R*; *Y*) increases monotonically with *p*^∗^(*y* = *r*|*r*).

### B.2 Softmax IB solution validations

Here we show that the softmax solution derived above is equal to the general IB solutions we present in Results for the tasks that we study. Namely, we validate that accuracy symmetry holds for each task and strategy we considered, then show that the softmax solution therefore minimizes the IB optimization problem. We also show that the softmax solution can hold when there are more than two latent states by examining a hypothetical three-state task.

To evaluate whether IB-optimal behavior followed (equation B5) in our tasks, we compared choice behavior produced using (equation B5) to IB-optimal behavior produced by the general IB optimization algorithm. As a strong test of (equation B5), we compared these two sets of solutions at matched values of *β* over the entire length of IB bound (i.e., using many different values of *β*). Specifically, we first computed solutions using (equation B5) over a large range of *β*^∗^ = *βα*^∗^ values. Then, we converted each *β*^∗^ to the corresponding *β* by computing *α*^∗^. These *β* values were then used to produce the equivalent set of solutions using the general IB optimization algorithm. Thus, (equation B5) is valid for a given IB bound only if the properties of *β*-matched solutions, like *I*(*X*; *R*) and *I*(*R*; *Y*), are exactly equal.

#### Main IB bounds for the bead-prediction and horse-prediction tasks

For the bead-prediction experiment, the fully optimal strategy was approximated using the previous seven beads to infer the identity of the hidden source jar. When we draw the IB curves for this strategy using (equation B5) and the IB optimization algorithm, they appear visually identical (Figure B14A).

Before taking a closer look, it is worth noting that the IB algorithm can be sensitive to the initialization point, especially for low or high values of *β*. When using a random initialization, the algorithm can sometimes converge to a suboptimal solution, making comparisons with the softmax solution difficult. We eliminated these problems by initializing the IB algorithm with the predicted softmax solution. We also confirmed that this approach did not bias the algorithm toward suboptimal solutions: for this bound (and for all other bounds we examined), softmax initialization did not differ from random initialization in terms of the eventual value of the IB cost function or of the value of *I*(*R*; *Y*) (as a function of *I*(*X*; *R*)), except for very small values of *β* (Figure B14B).

Because the task has a uniform *p*(*y*) by design and the solution computed using the general IB algorithm exhibits accuracy symmetry required of *p*^∗^(*r* = *y*|*y*) for all values of *β* (Figure B14C), the conditions required for the softmax solution to hold are met. Therefore, the softmax solution was identical to the general IB solution over the entire length of the IB bound (Figure B14D). *I*(*X*; *R*) and *I*(*R*; *Y*) are identical when the solutions are *β*-matched, validating the precise form of (equation B5). As a final sanity check, the maximum difference between the set of choice probabilities computed using the softmax solution and the one derived from the IB algorithm was effectively zero for each value of *β* (bottom plot).

The one-back heuristic IB bound for the bead-prediction task is also exactly equivalent to solutions provided by (equation B5), with no differences between the *β*-matched solutions (Figure B15).

For the three different sets of generative statistics (low, intermediate, and high W-S ratio) used in the horse-prediction experiments, the IB bounds for the fully optimal and equal-weights heuristic are also equivalent to solutions provided by (equation B5). For the sake of brevity, we show the analyses just for the intermediate W-S ratio conditions (Figures B16, B17). Readers interested in the analyses for all IB bounds (including the ignore-weak bounds) can find them in the publicly available analysis code.

#### IB bound for a task with three latent states

To demonstrate that (equation B5) can hold in cases with more than two latent states, we created and evaluated the fully optimal IB bound for a three-state version of the bead-prediction task. This hypothetical three-jar bead-prediction task featured three types of beads, such that each jar contained predominantly one unique bead type. The two less-common bead types for each jar were present in small, identical amounts. The transition structure was similarly symmetric: the stay probability was the same high probability for all jars, and the switch probability was the same low probability (details shown in Figure B18E). For this hypothetical three-state task, (equation B5) was equivalent to IB-optimal behavior for the fully optimal (seven-back) strategy (Figures B18).

**Fig. B14:**
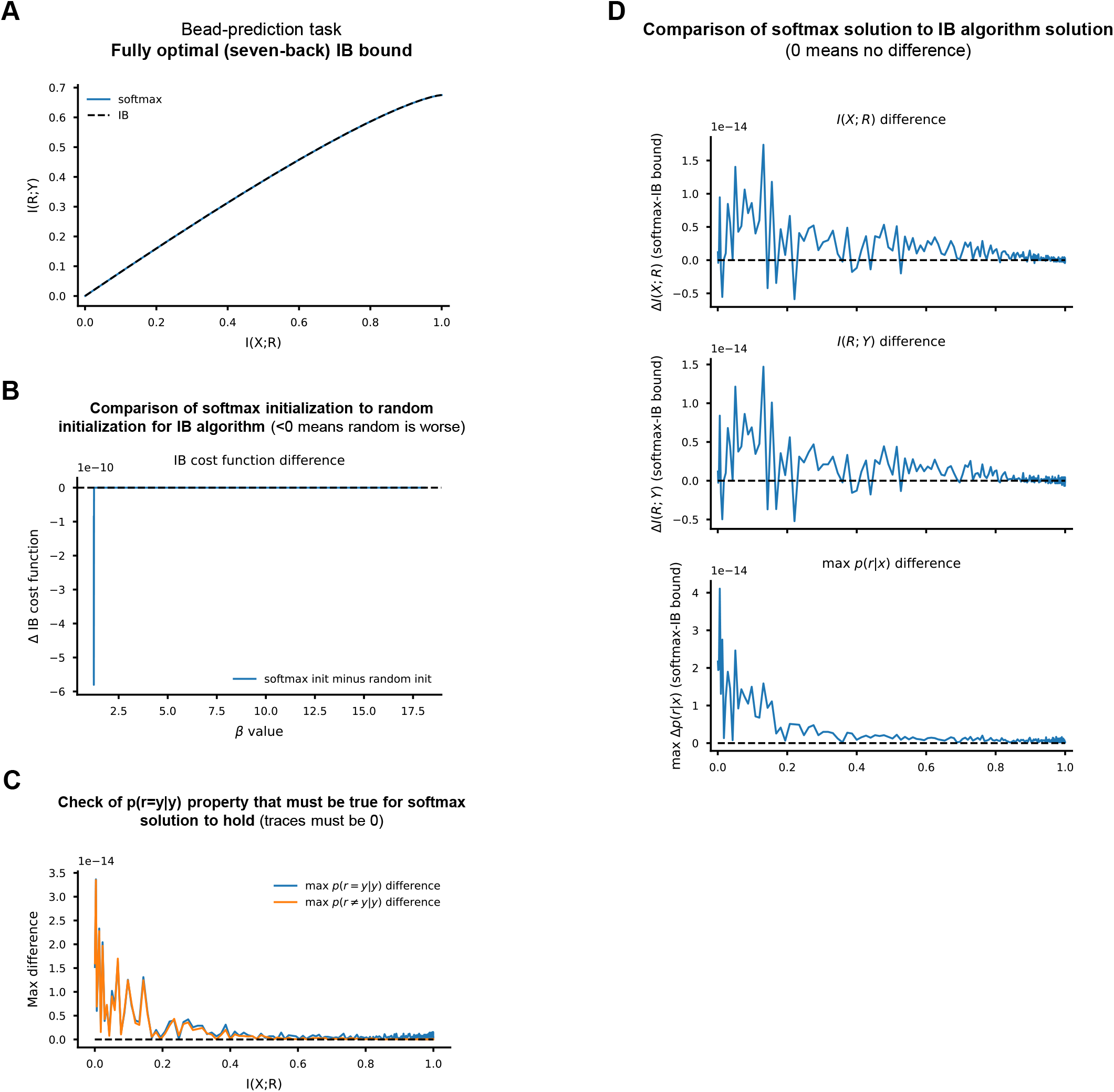
IB bound comparison for the fully optimal strategy for the bead-prediction task. **(A)** IB curves for the softmax solution ((equation B5)) and IB-optimal behavior produced using the general IB optimization algorithm (which completely obscures the softmax curve). **(B)** Difference in the IB cost function when initializing at the softmax solution minus random initialization, plotted as a function of *β*. Values *<* 0 indicate that random is worse. There was no difference in the value of the cost function between these two initialization schemes, except at low values of *β* where initializing at the softmax solution was better. **(C)** Test for accuracy symmetry required of *p*^∗^(*r* = *y*|*y*) for the softmax solution to hold. The blue trace gives the difference between the maximum and the minimum values of *p*^∗^(*r* = *y*|*y*) for all *y* ∈ *Y*. Values close to zero indicate that *p*^∗^(*r* = *y*|*y*) is the same for all *y* ∈ *Y*. The orange trace provides the same max-min difference value for the set of *p*^∗^(*r ≠ y*|*y*) for all possible values of *r* ∈ *R* and *y* ∈ *Y* (provided that *y ≠ r*). Both traces must be effectively zero (like they are here) for the required property to be present. **(D)** Comparison of *β*-matched solutions between (equation B5) and the general IB algorithm (using softmax solution initialization). Each plot shows the difference in a particular property, as indicated, between the softmax solution and the IB algorithm-derived solution as a function of *I*(*X*; *R*). Max *p*(*r*|*x*) deviation gives the maximum absolute difference across all values of *p*(*r*|*x*) between the two solutions. Note the ordinate scales in panels A–D (differences are tiny).

**Fig. B15:**
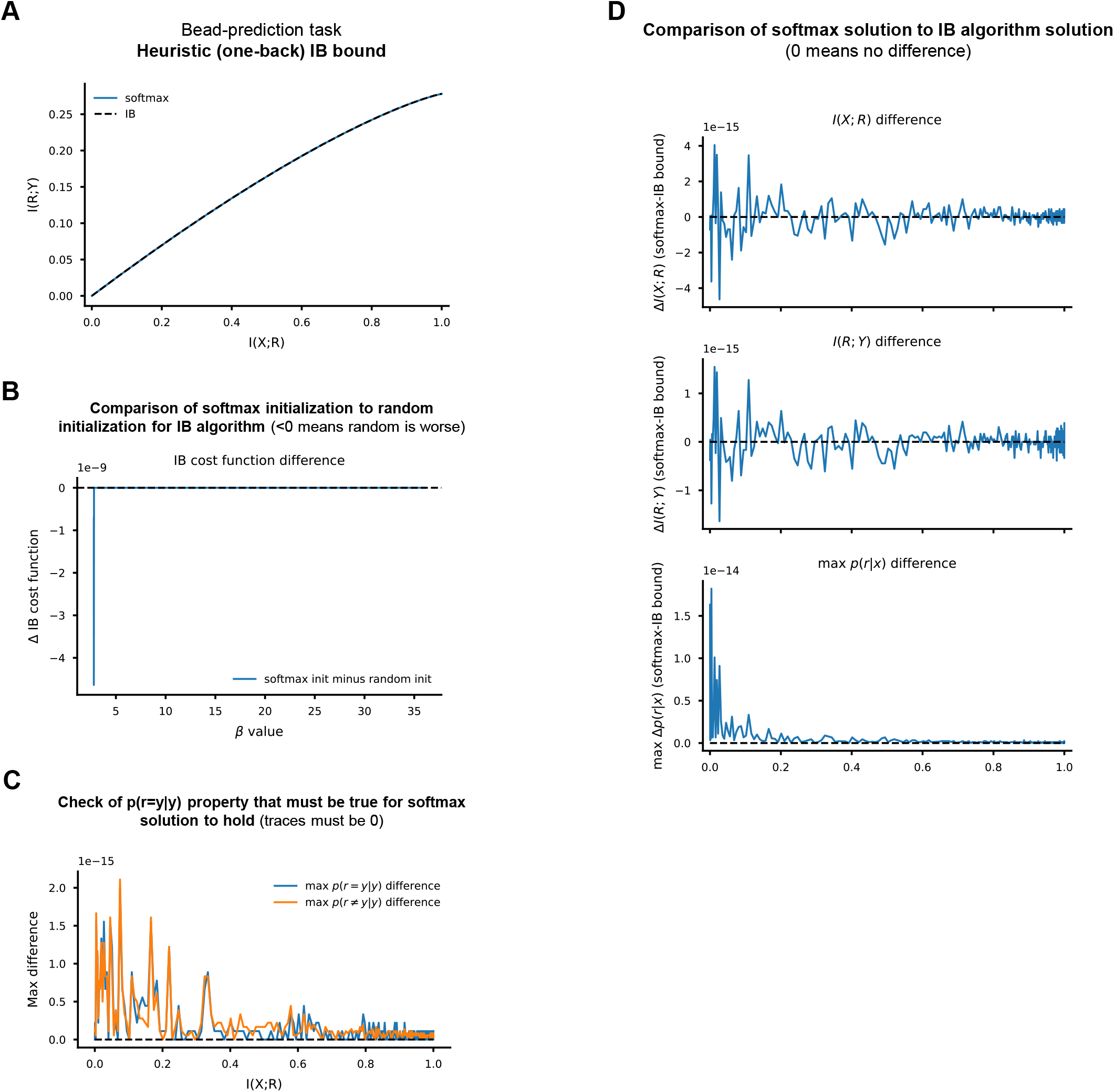
IB bound comparison for the one-back heuristic strategy in the bead-prediction task. **(A,B,C,D)** Same as for Figure B14, but for the one-back heuristic.

**Fig. B16:**
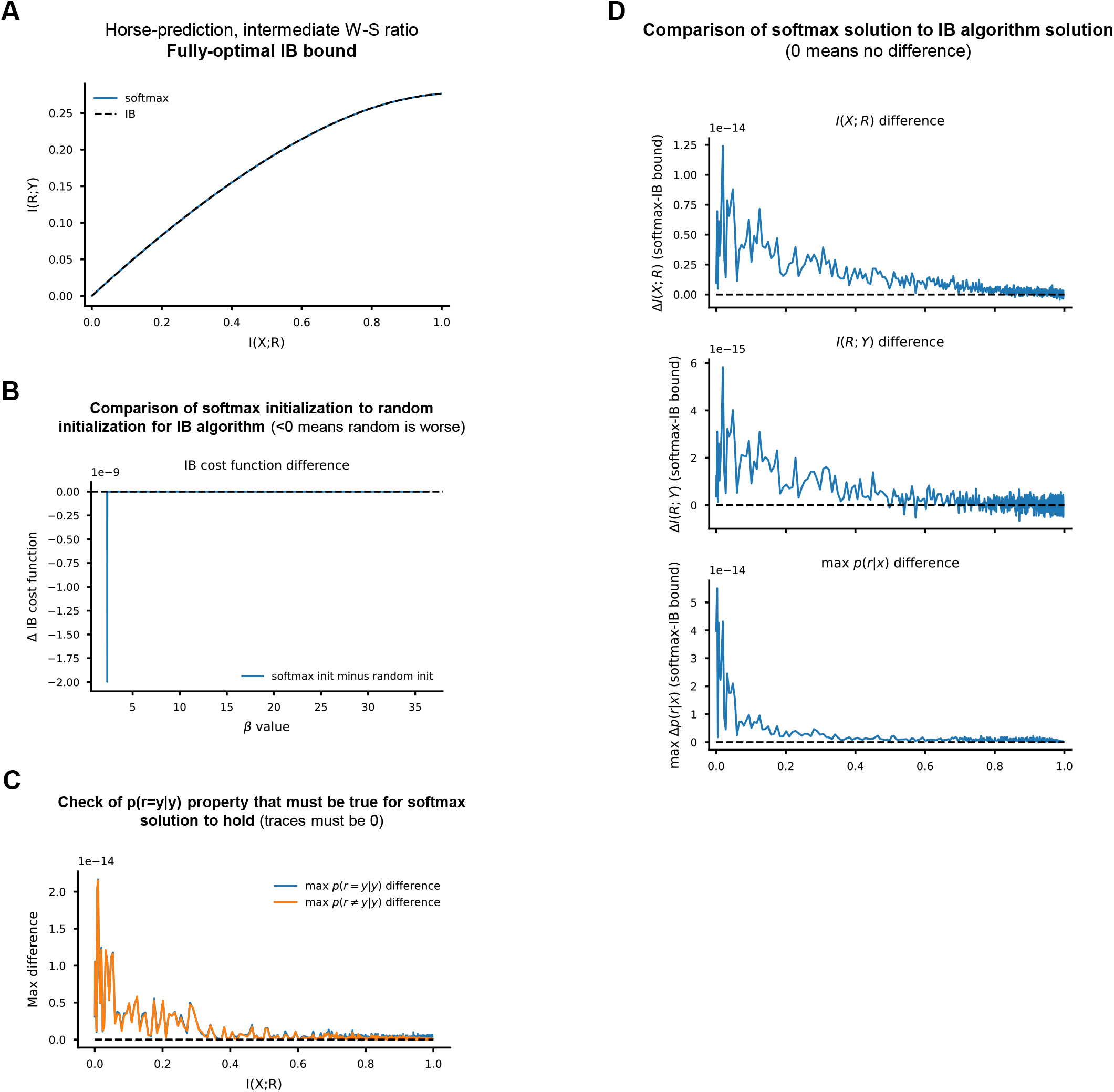
IB bound comparison for the fully optimal strategy in the horse-prediction task with the intermediate W-S ratio. **(A,B,C,D)** Same analyses as Figure B14.

**Fig. B17:**
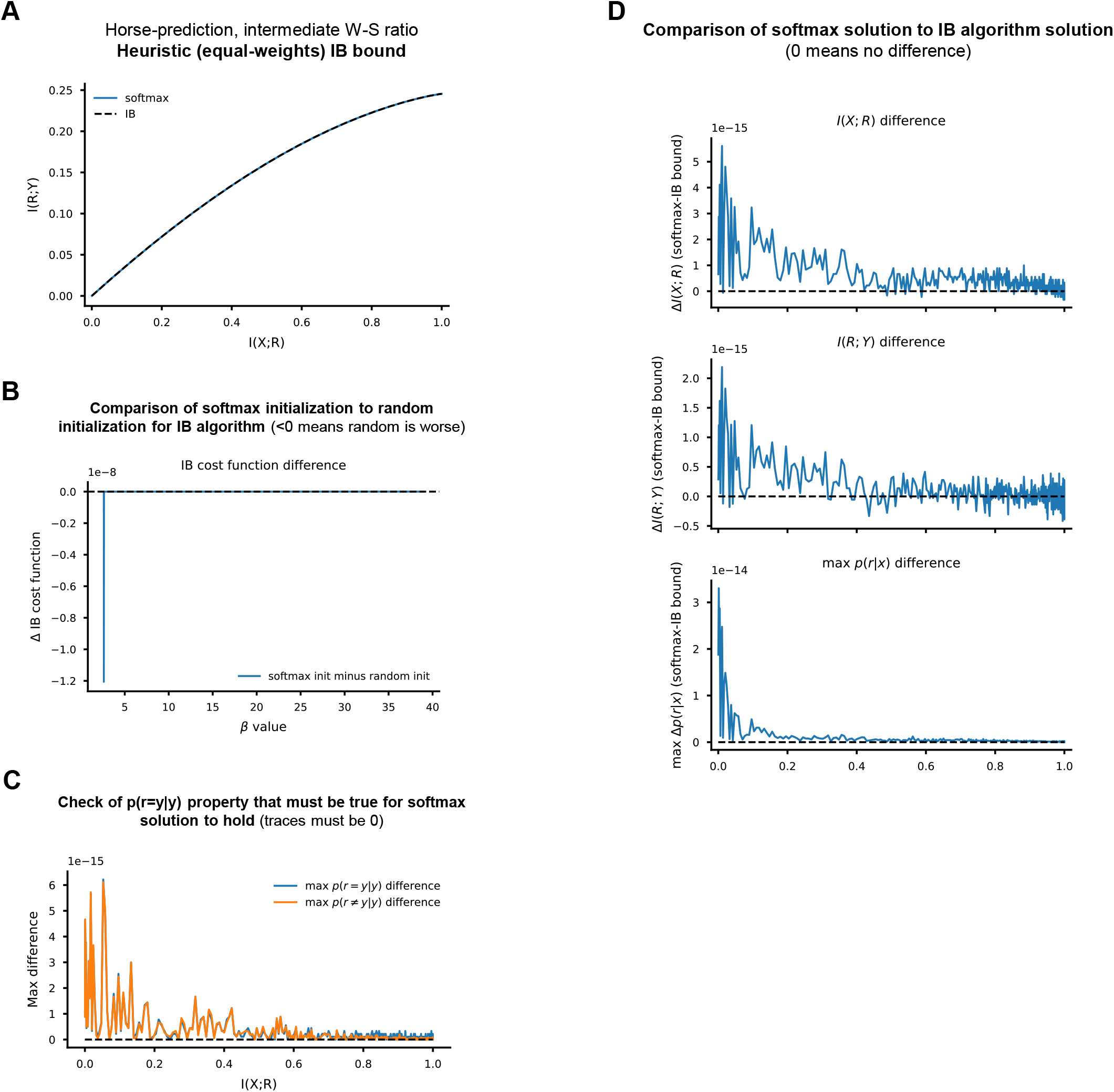
IB bound comparison for the equal-weights heuristic strategy in the horse-prediction task with the intermediate W-S ratio. **(A,B,C,D)** Same analyses as Figure B14.

**Fig. B18:**
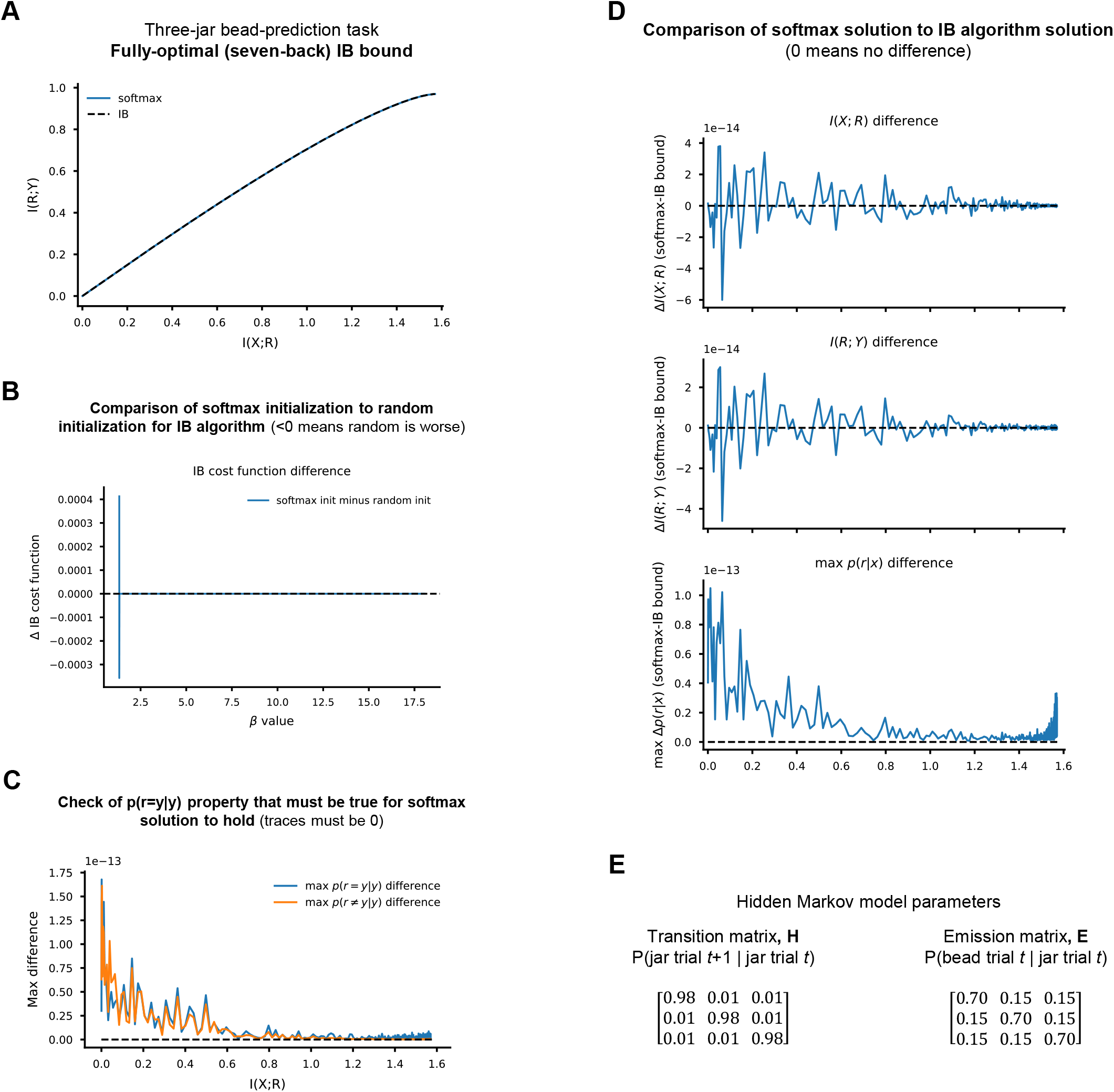
IB bound comparison for the fully optimal strategy in the three-jar bead-prediction task. **(A,B,C,D)** Same analyses as Figure B14. **(E)** The transition and emission matrices for the hidden Markov process representing this task.

